# Printing double network tough hydrogels using Temperature-Controlled Projection Stereolithography (TOPS)

**DOI:** 10.1101/2023.03.20.533294

**Authors:** Puskal Kunwar, Bianca Louise Andrada, Arun Poudel, Zheng Xiong, Ujjwal Aryal, Zachary J Geffert, Sajag Poudel, Daniel Fougnier, Ivan Gitsov, Pranav Soman

**Affiliations:** Syracuse University, Biomedical and Chemical Engineering Department; BioInspired Institute, Syracuse, New York, 13210, USA; State University of New York ESF, Department of Chemistry, Syracuse, New York, 13210, USA; The Michael M. Szwarc Polymer Research Institute, Syracuse, New York, 13210, USA

**Keywords:** double network hydrogel, projection stereolithography, additive manufacturing, digital micromirror, mechanically reconfigurable soft devices

## Abstract

We report a new method to shape double-network (DN) hydrogels into customized microscale 3D structures that exhibit superior mechanical properties in both tension and compression. A one-pot prepolymer formulation containing photo-cross-linkable acrylamide and thermo-reversible sol-gel κ-carrageenan with a suitable crosslinker, and photo-initiator/absorbers are optimized. A new TOPS system is utilized to photo-polymerize the primary acrylamide network into a 3D structure above the sol-gel transition of κ-carrageenan (80°C), while cooling down generates the secondary physical κ-carrageenan network to realize tough DN hydrogel structures. 3D structures, printed with high lateral (37μm) and vertical (180μm) resolutions and superior 3D design freedoms (internal voids), exhibit ultimate stress and strain of 200 kPa and 2400% respectively under tension, and simultaneously exhibit high compression stress of 15 MPa with a strain of 95%, both with high recovery rates. The roles of swelling, necking, self-healing, cyclic loading, dehydration, and rehydration on the mechanical properties of printed structures are also investigated. To demonstrate the potential of this technology to make mechanically reconfigurable flexible devices, we print an axicon lens and show that a Bessel beam can be dynamically tuned via user-defined tensile stretching of the device. This technique can be broadly applied to other hydrogels to make novel smart multifunctional devices for a range of applications.

## 1. Introduction

Hydrogel materials have found applications in drug delivery, tissue engineering, biosensing, soft robotics, flexible electronics, and soft photonics. However, traditional single-network hydrogels typically exhibit inferior mechanical properties which limit their use in the field.^[1–5]^To address this challenge, double network (DN) hydrogels have been developed for applications that require superior toughness, stretchability, and compressive strength.^[6–13]^ DN hydrogels typically consist of two entangled networks that can be polymerized using two independent stimuli; one network allows energy dissipation during deformation while the other network provides toughness and/or stretchability.^[6,11–17]^ Many one-pot synthesis strategies (sol-gel transitions, click chemistries, sequential polymerization) have been used to synthesize DN gels, however, shaping them into customized 3D structures remains a significant challenge.^[7–9,17–24]^Conventional molding and casting methods are used to generate simple geometries such as sheets, slabs, and discs, ^[8,25,26]^ while extrusion-based 3D printing have also been used to print customized 3D shapes, although at low resolution and speeds.^[7,27–29]^ Light-based printing methods can print at high-resolutions, the ability to rapidly photo-crosslinking with DN hydrogels remains a significant materails challenge.^[20,30–33]^

Bottom-up Projection Stereolithography (PSLA) has emerged as the favorite light-based 3D printing method due to its capability to make customized parts with microscale resolution and superior design flexibility.^[34–37]^ A typical setup consists of spatially modulated light patterns projected through a transparent bottom window to crosslink photosensitive liquid resin in XY plane before moving the stage up (Z-direction) to print the structure in a layer-by-layer or layerless continous manner.^[34,38,39]^ Unfortunately, many DN resins do not meet the criteria of low viscosity and rapid photo-crosslinking at specific wavelengths. For instance, thermos-reversible sol–gel transitions require elevated temperatures beyond the operating range of current printers, while reaction durations and crosslinking times for orthogonal click chemistries remain too long for PSLA (hours).^[40–42]^ New strategies that combine PSLA technology with DN hydrogels could potentially lead novel soft devices with superior mechanical properties.

Here, we report a simple one-pot PSLA printing strategy that combines the advantages of PSLA (rapid, high-resolution, 3D design flexibility) and hydrogels (transparency, hydration) to shape DN gels into complex geometries with superior mechanical properties (high strength in both tension and compression). To demonstrate how this strategy can be used to make multifunctional soft devices, we printed an axicon lens using DN gel and showed that dynamic stretching can be used to modulate its optical performance. This work can potentially expand the design freedoms and the material library for making stimuli-responsive soft devices using DN hydrogels.

## 2. Results

### 2.1. TOPS design and printing of DN hydrogels

Single pot DN hydrogel prepolymer solution consists of acrylamide monomers, κ-carrageenan, crosslinker MBAA, LAP photoinitiator, and in some cases a photo-absorber. The basic mechanism of DN formation involves photo-crosslinking of the acrylamide network and physical crosslinking of κ-carrageenan below its sol-gel transition temperature (80°C) (**Figure 1A**).^[43,44]^ Thus, a new stage was designed and built to allow the printing of the prepolymer formulation above 80°C. Below 80°C, prepolymer physically crosslinks into a viscous gel and cannot be printed using PSLA. TOPS setup consists of a light source, diffuser, DMD, projection optics, z-stage, and a specifically designed sample holder for maintaining a constant elevated temperature. (**Figure 1B**). The design of the sample holder consisted of a copper plate with a center hole embedded inside the PDMS bath. The 16 mm hole serves as the fabrication window and the copper plate heats the sample holder. A multistep molding and casting technique was used to build the sample holder.

**Figure 1.**
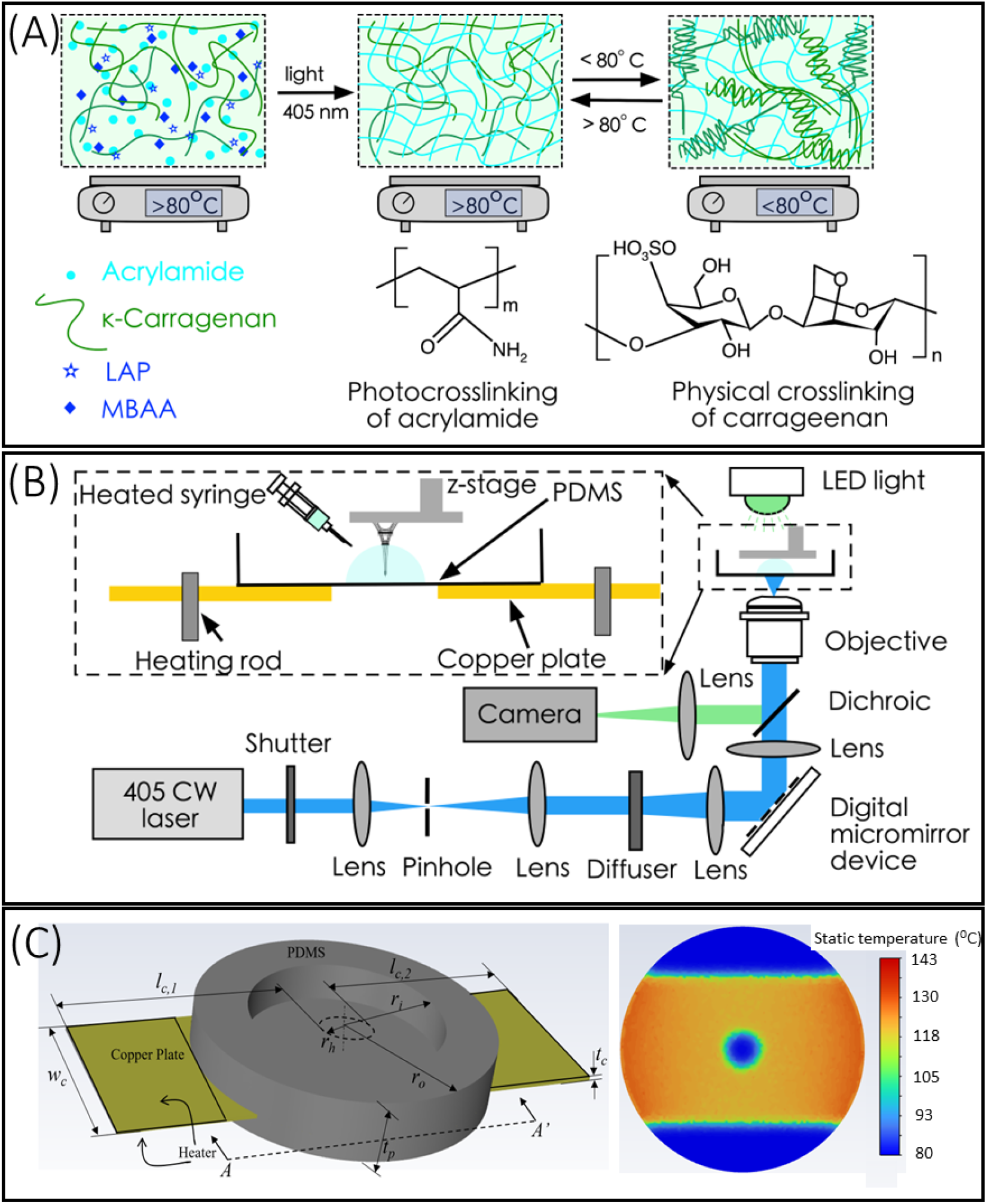
(A) Schematic diagram of DN gels formed by photocrosslinking and thermos-reversible sol-gel transition. Acrylamide in the presence of crosslinker MBAA and photoinitiator LAP formed the first network through irreversible photocrosslinking. κ-carrageenan formed the second network by reversible physical crosslinking via sol-gel transition (B) Schematic setup of the TOPS hydrogel for printing 2D and 3D DN gels structures. (C) The 3D model of the sample holder was used as a computational domain consisting of a PDMS dish with a copper plate. Temperature distribution over the PDMS layer at the plane corresponding to section A-A’ at t=90 sec when the steady state was reached.

Computational fluid dynamics (CFD) simulation provided the temperature distribution to guide the design of the sample holder (**Figure 1C, S1-S3**). The temperature within the fabrication window was experimentally measured to be 80°C when the heaters were operated at 150°C (423 K). DN hydrogel structures were printed using PSLA at 80°C as explained below. Based on user-defined CAD design, spatially modulated light patterns are irradiated onto the liquid prepolymer solution maintained at an elevated temperature. Upon irradiation, the LAP photoinitiator absorbs light and initiates a polymerization reaction in the presence of a crosslinker (MBAA) to form a polyacrylamide network that locks-in place target geometry followed by cooling-driven physical crosslinking of the κ-carrageenan network resulting in a DN hydrogel structure. For each layer, prepolymer solution was injected via a heated syringe into the fabrication window area, and post-printed samples were developed in hot DI water (at 80°C) for 2 mins to remove any uncrosslinked monomers.

### 2.2. Printing 2D, 3D solid, and 3D hollow DN structures at microscale resolution

Before printing complex, 2D/3D structures using DN hydrogels, both lateral and vertical resolution limits were characterized. For quantifying the lateral resolution of TOPS, digital masks of intersecting line patterns with varying pixels (1-10) were printed using a laser intensity of 2.17mW/cm^2^ and an exposure time of 15 seconds. Measured line widths were close to the theoretical resolution, which corresponds to the micromirror size in the Digital Micromirror Device (DMD) chip in PSLA (**Figure 2A-B**). For instance, the smallest feature size of 37 μm was experimentally obtained for a 3-pixel line pattern, close to the theoretical resolution of 36 μm. The vertical printing resolution, necessary to generate structures with internal voids, was characterized by adding a photo-absorber (tartrazine, 0.006 wt%) to the prepolymer solution. The curing depth was optimized by varying the exposure dose, a function of light intensity and time. Here, we printed a rectangular slab on the coverslip by varying the exposure time while maintaining a constant laser intensity of 2.4 mW/cm^2^. The thickness of the structures was measured to obtain the curing depth and plotted as a function of exposure time (**Figure 2C**). A z-resolution of 180 μm was obtained for the exposure time of 15 seconds. The curing depth can be tuned from 180 μm to 420 μm by varying the exposure time from 15 to 60 seconds. Exposure time below 15 seconds did not result in crosslinking of DN gels structures.

**Figure 2.**
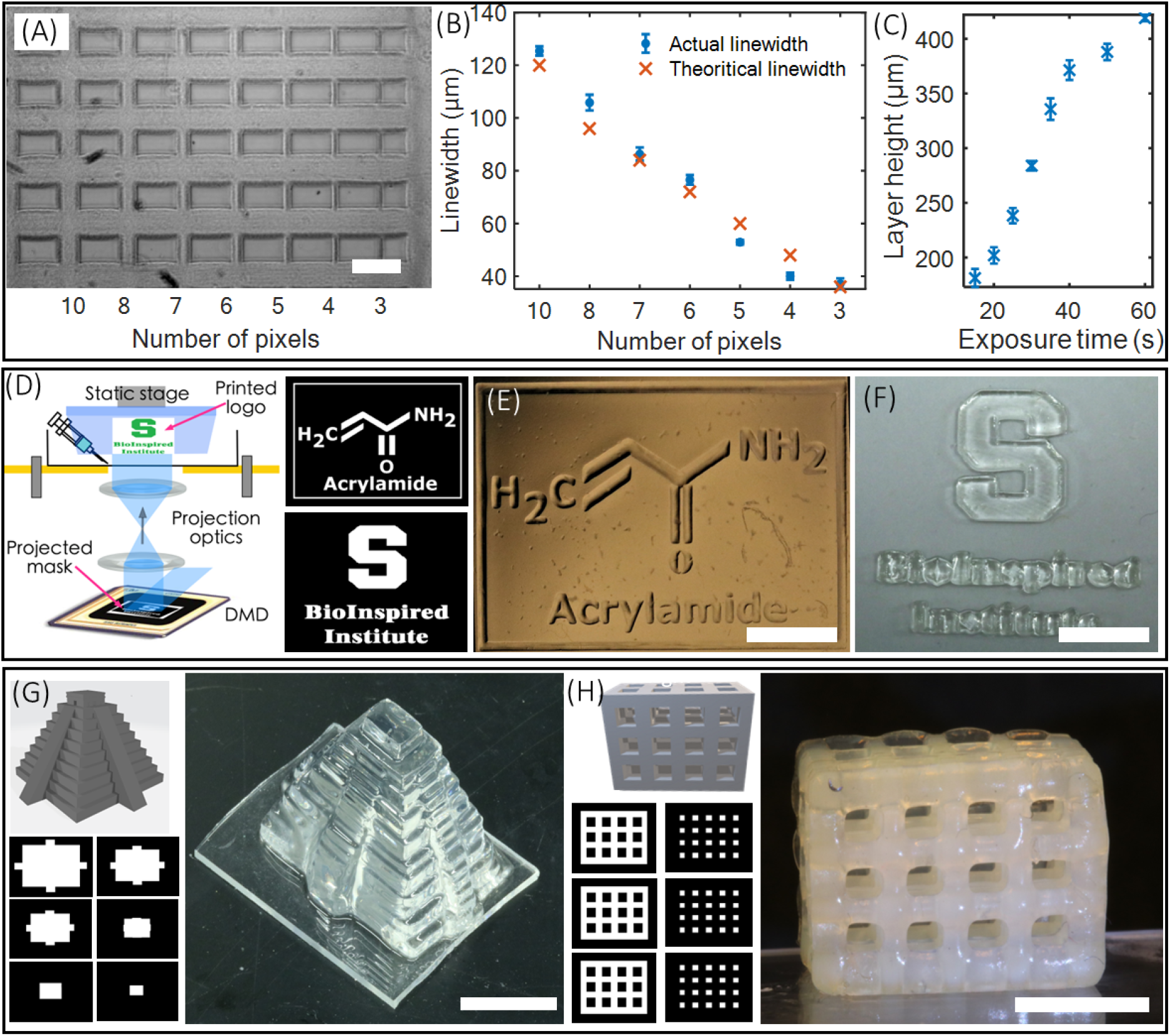
(A-B) Figure and plot depicting the lateral resolution of printed structure using DN gels (scale bar-200 μm). (C) Plot showing the axial resolution of printed DN gels structure. (D) Schematic of 2D printing of planar structure and computer-generated digital mask for printing chemical structure of acrylamide and logo of Bioinspired Institute. (E-F) 2D printed acrylamide structure before the development and Logo of Bioinspired Institute after the development of structure (scale bar-4 mm). (G) 3D CAD model, corresponding computer-generated digital mask and 3D printed structure of Mayan pyramid. (scale bar-5mm). (H) CAD model, digital masks, and 3D printed lattice structure. The printed structure was dipped into ethanol for 30 minutes to remove tartrazine and enhance the contrast for imaging (Scale bar-5mm).

Results show a 2D pattern of the chemical structure of “acrylamide” and the logo of the Bioinspired Institute of Syracuse University (**Figure 2D-F**). Next, a solid 3D geometry in form of a Mayan pyramid was printed with a laser intensity of 2.17 mW/cm^2^ and a layer exposure time of 15 seconds per layer (**Figure 2G**). Lastly, hollow 3D geometry with overhangs and undercuts was chosen and printed (**Figure 2H**). Here, a 0.006 wt% tartrazine photo absorber was added to the prepolymer solution before printing the geometry (2.4 mW/cm^2^, exposure time per layer = 15 seconds). All printed structures were developed in water (80°C for 2 minutes) to remove uncrosslinked monomers.

### 2.3 Tensile performance of printed DN gels structures

Tensile properties of TOPS-printed DN dog-bone shaped structures (**Figure 3A(i), Video V1**) were compared with identical structures made from single-network structures (acrylamide-only, κ-carrageenan only). Acrylamide-only dog-bones structures were printed with TOPS (2.17 mW/cm^2^) using single exposure while conventional molding and casting was used to generate dog-bone geometry using κ-carrageenan. The representative stress-strain plot shows the superior fracture energy (1238.1 J/m^2^) of DN structures as compared to single-networked acrylamide (425.9 J/m^2^) and κ-carrageenan (4.25 J/m^2^) structures (**Figure 3A(iii)**). The ultimate stress required to stretch the hybrid gel structure by 10.6±1.64 times was 175±37 kPa, whereas the stress and associated ultimate strain for acrylamide gels were 28±3.6 kPa and 23.5±3.7 kPa (**Figure 3A(iii)**). The κ-carrageenan structure breaks at the strain of 0.8±0.06 at the stress of 10.5±4.9 kPa (**Figure 3A(iii)**). The modulus of elasticity is 79.5±22 kPa highest for hybrid gel whereas it was 8.2±0.72 kPa for acrylamide and 7±1.4 kPa for κ-carrageenan. Next, hollow-lattice geometry with a strut width of 900 μm was printed at 2.4 mW/cm^2^ with an exposure time of 15 sec per layer (**Figure 3B**). This structure can withstand a load of 75grams for 20 seconds by stretching 8 times its original length (**Figure 3B, Video V2**). Overall, these results indicate that the hybrid gel structure obtains its strain from the polyacrylamide polymer, and the addition of κ-carrageenan increases the stiffness properties. It is suggested that a double helical structure in the κ-carrageenan starts to break at a small strain that unzipped progressively as the strain increases and leads to permanent deformation. The breaking of cross-linked double helices of κ-carrageenan serves as a sacrificial bond, which greatly helps to improve the stiffness of DN hydrogel through energy dissipation.^[43]^Throughout the process, the acrylamide network remains intact and maintains the geometry of the printed structure. Standard dog-bone geometry was used to characterize the influence of many processing variables on tensile regimes.

**Figure 3.**
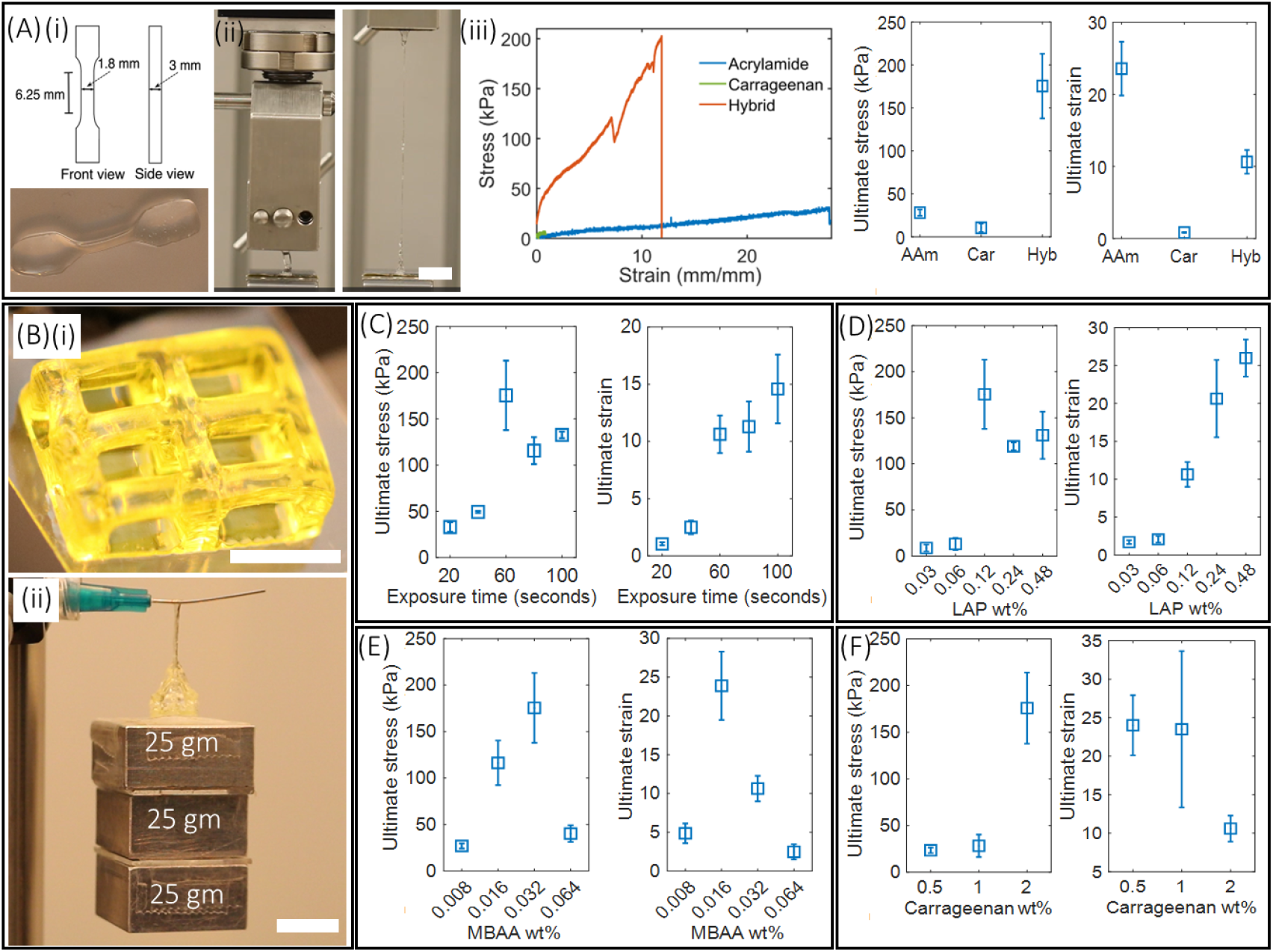
(A)(i) Schematic and printed dog-bone structure using a hybrid gel structure with laser power of 2.17 mW/cm^2^ and exposure time of 60 seconds. (ii) Photographs showing the performance of dog-bone structure during tensile testing and the structure stretched 9.5 times its original length. (iii) Stress-strain plot obtained from dog-bone structures printed using DN gel, acrylamide-only gel, and carrageenan-only gel. Ultimate stress and ultimate strain of the fabricated structures are also depicted. (B) (i) 3D printed hollow-lattice geometry with a structure for studying the tensile performance. (scale bar– 5mm). (ii) photographs showing the tensile performance of the structure (scale bar-8 mm). (C-F) Ultimate stress and ultimate strain of DN gel structures printed by varying exposure times, and amounts of photoinitiator, cross-linker (MBAA), and κ-carrageenan.

### 2.4 Influence of DN formulations on tensile properties

#### 2.4.1 Role of light dosage

Dogbone DN gels structures were printed using TOPS under single exposure conditions (in this case, the z-stage does not move) by varying the exposure times from 20 to 100sec at a constant laser intensity of 2.17 mW/cm^2^. Structures printed using the exposure time of 40 seconds and below broke with a very small stress of 50 kPa and exhibited a strain of less than 3 (**Figure 3C**). The structures printed with exposure times of the 20s and 40s were underexposed and the associated covalent network of acrylamide is partially formed so that they exhibit less stretchability and accompanying stress. The highest ultimate stress of 175±37 kPa was found for the exposure time of 60 sec. Above this exposure time of 60 sec, there is a slight decrease in ultimate stress, but the difference is not significant (Figure 3C). Similarly, the difference in strain is also minimum among the structure printed above 60 sec although the highest ultimate strain of 14.59±3 is observed for the structures printed with the longest exposure time of 100 sec (**Figure 3C**). Further, the elastic modulus remains almost same for the structures printed with different exposure times (**Figure S4**). Representative stress-strain plots obtained from the hybrid gel structures irradiated with different exposure times and the corresponding elastic modulus obtained are depicted in SI (**Figure S4**).

#### 2.4.2. Role of photoinitiator concentration

Dogbone DN samples were fabricated at a laser intensity of 2.17 mW/cm^2^ and an exposure time of 60 seconds using varying concentrations of LAP (0.03wt% to 0.48 wt%). Structures with 0.03 wt% and 0.06 wt% LAP did not perform well while stretching and exhibited ultimate stress of less than 13 kPa (**Figure 3D**). The stress increased to 175.3±37 kPa for the structure printed with 0.12 wt% LAP. Further, an increase in the concentration of the photoinitiator to 0.24 wt% and 0.48wt% decreased the ultimate stress to 119±4 kPa and 131±25 kPa (**Figure S5**). There was an increasing trend in terms of ultimate strain and the maximum strain of 25.99±2 kPa for printed structures with the highest LAP concentration (**Figure 3D**). Representative stress-strain plots obtained from the hybrid gel structures printed by varying photoinitiator concentration and corresponding variation in the modulus of the structure are reported in SI (**Figure S5**). The modulus reaches a maximum of 79±22 kPa for 0.12wt% and dropped to 28±5.4 kPa and 33.8±4.6 kPa for 0.24 wt% and 0.48 wt% (**Figure S5**).

#### 2.4.3. Role of crosslinker concentration

Dogbone DN samples were printed using TOPS with varying concentrations of MBAA (0.008wt% to 0.128wt%). Results showed that the highest ultimate stress of 175±37 kPa and the ultimate strain of 10±1.64 kPa was observed for the 0.032 wt% MBAA. The ultimate stress decreased to 116±24 kPa while the ultimate strain increased to 23.8±4.4 for the MBAA concentration of 0.016 wt% (**Figure 3E**). These parameters decreased significantly when the MBAA concentration decreased to 0.008 wt%. Furthermore, an increase in the concentration of MBAA crosslinker to 0.064 wt% and 0.128 wt % decreased both the stretchability and force required to break the dog-bone structure. The structure printed using an MBAA concentration of 0.128 wt% readily broke, and hence was omitted for further studies (**Figure 3E**). The elastic modulus of 79.5±22 was highest for an MBAA concentration of 0.032 wt% and the moduli were less than 30 kPa for structures printed with the other MBAA concentrations (**Figure S6**). Results show that an increase in the concentration of crosslinker increases the crosslinking density, which leads to a short polyacrylamide chain and low fracture energy. For very high or low MBAA concentrations, the covalent network becomes too complaint, leading to deformation of the network with small stress.

#### 2.4.4. Role of κ-carrageenan concentration

Dogbone DN samples were printed with three concentrations of κ-carrageenan: 0.5 wt%, 1 wt%, and 2 wt% as above this concentration, κ-carrageenan did not dissolve in the water. As expected, the lower concentration κ-carrageenan structures were soft and highly stretchable. The strain decreased from 24±3.9 to 10.6±1.69, when the concentration of κ-carrageenan increased from 0.5 wt % to 2 wt%, whereas the ultimate stress increased from 23.5±3 kPa to 175±3 kPa (**Figure 3F**). The elastic modulus of 79.5±22 kPa was highest for the structure printed with 2 wt%κ-carrageenan and decreased with decreasing κ-carrageenan concentration (**Figure S7**). This result suggests that the κ-carrageenan increases the stiffness properties whereas the acrylamide contributes to the stretching properties of the structure printed with hybrid gels.

### 2.5 Necking and solution to necking phenomena

Many studies on DN hydrogels report a necking phenomena, although they failed to offer a solution to this behavior.^[43,45,46]^ We also observe similar necking behavior with printed DN structures at strain of ~1.5 with variations in both their numbers and locations. Necking location corresponded to the point of breakage when more strain was applied. Furthermore, once necking is initated, sample were not able to recover back to their original size. (plastic deformation occurs) (**Video V1, Figure 4A**). It has been proposed that necking occurs due to spatial variability within the material structure, which causes the material to experience disproportionate stress during deformation, leading to instability and resulting in localized strain in a specific area.^[47]^ We hypothesize that the cause of necking in printed DN samples is due to inhomogeneous spatial distribution of water within the structure, as the surface of the printed structure loses water faster than the center of the structure during the printing and post-processing steps. To test this hypothesis, we immersed printed DN samples in water for 5min prior to recording stress-strain plots. Plots and video show that the structures did not show the necking phenomenon (**Figure 4A(i-ii), Video V1**), but exhibited lower stress compared to the as-printed structure possibly due to an increase in the total water content from 82% to 87%. To test this, another prepolymer solution was used to print dogbone sample with an initial (as-printed) water content of 60%, and then immersed in water for 15min to get the total water content to 82% before tensile testing. Results show a recovery of stiffness and associated stress with no necking behavior. (<u>Green curve</u>, depicted as <u>restored</u> in **Figure 4A (i)**) A detailed study of the swelling of the structure and the influence of swelling on the tensile properties of the structure is mentioned in SI (**Figure S8-S11**).

**Figure 4.**
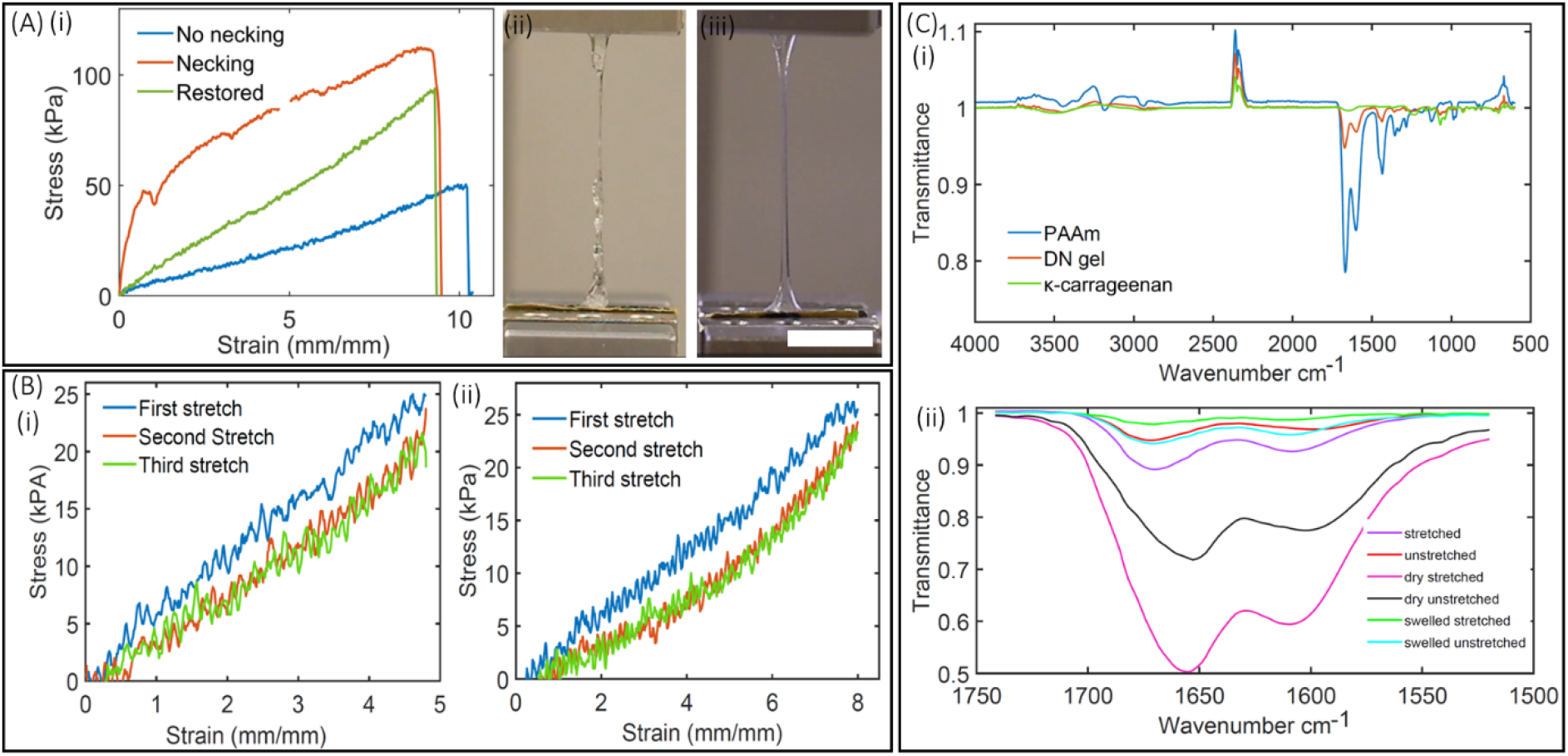
(A) Solution to the necking behavior (i) Comparison of the stress-strain plot of structure with necking behavior (red), without necking behavior (blue), and without necking with restored stress (green) (ii-iii) photographs showing the necked and un-necked structure during stretching. (B) Stress plotted during three cycles of loading and reloading (i) with the strain of (480%) (ii) with the strain of (800%) (C) (i) FTIR spectra of molded and casted κ-carrageenan, TOPS-printed PAAm, and DN gels (ii) FTIR spectra, zoomed in, acquired from as-printed, dry, and swollen DN gels structures before and after stretching.

### 2.6 Response to cyclic loading-unloading and FTIR studies of DN gels

#### 2.6.1. Cyclic loading-unloading

DN dog-bones, hydrated for 10 minutes, were stretched to the strain of 480% for three repetitions, and fracture energy was calculated for all the repetitions. Results showed that the fracture energy for the first cycle was 61.059 J/m^2^, which decreased to 44.92 J/m^2^ for the second cycle, however, the loss was only 0.94 J/m^2^ from the second to the third cycle (**Figure 4 B(i)**). Similarly, stretching the structures to the strain of 800% for the three-repetition cycle showed a decrease in fracture energy from 98.67 J/m^2^ to 69.41 J/m^2^ during the first and the second cycle while minimal energy loss of 1.93 J/m^2^ from the second to the third cycle (**Figure 4 B(ii)**). Overall, energy loss increases with increasing strain (from 480% to 800%).

#### 2.6.2. FTIR studies of DN gel structure

FTIR spectra of casted κ-carrageenan, TOPS-printed PAAm, and DN gels (dried, and swollen) are shown and the FTIR fingerprints associated with functional groups are assigned (**Figure 4C(i)**, Table S2). FTIR spectra were acquired from as-printed, dry, and swollen dog-bone DN gel structures before and after tensile loading. The dry samples were obtained by drying them overnight at room temperature and the swollen sample were obtained by immersing them in water for 4 hours. Of particular interest in this study are bands corresponding to moieties that participate in hydrogen bonds, such as the amide I (C=O stretching) band and amide II (N–H bending) band (**Figure 4C(ii)**). The observed FTIR bands and their assignments are summarized in Table 1. The amide I stretching band responds to the extent of hydration of the DN gel sample due to changes in the degree of hydrogen bonding. This band is shifted to a lower wavenumber in dried samples compared to hydrated as-printed and swollen samples due to an increase in intermolecular polymer-polymer hydrogen bonding. As evidenced by the absence of band shifting after tensile loading, the amide I band is relatively insensitive to changes in inter- and intra-molecular forces induced by mechanical loading.^[48]^

**Table 1.** FTIR fingerprints associated with functional groups amide I (C=O stretching) band and amide II (N–H bending) band obtained from polyacrylamide, as-printed, dry, and swollen DN gels dog-bone structures. ν stretching; δ, bending.^[48]^

| PAAm Reported | PAAm | DN gel structure | DN gel structure (Stretched) | Dried DN gel structure | Dried DN gel structure (Stretched) | Swollen DN gel structure | Swollen DN gel structure (Stretched) | Assignment |
| --- | --- | --- | --- | --- | --- | --- | --- | --- |
| 1618 | 1600 | 1598 | 1608 | 1602 | 1610 | 1609 | 1609 | $\delta$, NH<sub>2</sub> amide II |
| 1660 | 1668 | 1672 | 1670 | 1653 | 1655 | 1671 | 1671 | $\nu$, C=O amide I |

The amide II bending band is affected by tensile loading in as-printed and dried samples. Wavenumbers increase by 8-10cm^-1^ to approximately 1609cm^-1^ following tensile loading as compared to the equivalent unloaded control observed at ~1598cm^-1^. This is indicative of a reduction in inter- and intra-molecular interactions between polymer chains due to mechanically induced physical separation of the polymer chains. This can be attributed to the fracture energy loss during the structure’s cyclic loading (**Figure 4B**). Notably, no such mechanically-induced band shift was observed for swollen samples, where the amide II band appears at 1609 cm^-1^ both before and after loading. The main reason for this phenomenon is the complete solvation of the polymer matrix with hydration shells forming around the polymer chains. The net result is a marked reduction in polymer-polymer interaction in the fully swollen state. ^[48]^

### 2.7 Self-healing behavior

To assess if the thermoreversible sol–gel transition of the κ-carrageenan network can exhibit healing behavior at 80°C, a pyramid-shaped structure was 3D printed and cut into half using a sharp razor blade and was stained with pink and blue dye. The two-piece of the structure was sealed in a polyethylene bag and stored in a water bath of 85 °C for 20 minutes. The two halves of the structure self-heal to produce a complete pyramid (**Figure S12 A,B**). To quantify the self-healing behavior, a rectangular slab geometry, printed using TOPS, was cut in half using a sharp razor blade. To visualize the self-healing interface, one of the halves was stained with faint blue dye. Then, the two-piece was placed in close contact and heated to 80 °C for 20 minutes, and cooled down to initiate physical crosslinking of the κ-carrageenan network (**Figure S12 C,D**). The self-healed monolithic structure was able to withstand a weight of 70 gm although the self-healed interface starts to rupture when the load is increased to 200 gm (**Figure S12 E,F**). Similar experiments with self-healed DN dogbone samples resulted in an ultimate stress of 24±3 kPa and breaks at a strain of 2.5±0.6, much lower than as-printed samples (ultimate stress of 125 kPa and ultimate strain of 15.1) **(Figure S12 G).**This is because self-healing takes place solely via physical crosslinking of the κ-carrageenan network.

To explain the self-healing behavior, thermogravimetric analysis of dehydrated DN gel samples (n=4) was performed. Results showed a three-stage thermal decomposition (**Figure S13**). The onset of the first stage occurred at 224.10±3.08°C with a peak rate of mass loss at 236.03±1.38°C. The second stage had an onset of 263.99±0.80°C and reached a peak rate of mass loss at 279.83 ±0.54°C. The third stage had an onset of 360.37 ±0.82°C and reached a peak rate of mass loss at 382.04 ±0.91°C. The residue weight percent at 600°C was 25.00± 0.75%. As shown, the onset of thermal decomposition is well in excess of the thermal processing conditions utilized in both printing DN gel structures and inducing self-healing of the physically crosslinked network. Though TGA was performed on dehydrated samples, these results are believed to be representative of the thermal stability of hydrated DN gel structures due to the absence of hydrolysis-sensitive linkages at neutral pH. The self-healing behavior exhibited by DN gels can therefore be ascribed to the interdiffusion of solvated κ-carrageenan chains and subsequent reformation of physical crosslinks rather than decomposition into highly adhesive oligomers.

### 2.8 Compressive performance of DN hydrogel structure

Mechanical properties of DN cylindrical stub structures (radius=7mm and height=5mm) under compression were compared with identical structures made from single-network structures (acrylamide-only, κ-carrageenan only). Cylindrical stubs, printed using 2.17 mW/cm^2^ and an exposure time of 70 sec, were subjected to uniaxial compression, and associated stress and strains were plotted (**Figure 5A**). Results show that the ultimate compressive stress in the case of the DN structure was 15 MPa at a strain of 95%; 10 and 150 folds the stress possible by acrylamide and κ-carrageenan structures respectively. κ-carrageenan-only structures fracture at 0.032 MPa and a strain of 50% while polyacrylamide-only structures fracture at 1.45 MPa and a strain of 89% (**Figure 5A**). The result suggests both the κ-carrageenan network and acrylamide network increase the toughness of the hybrid structure not just by simple interpenetration but also through a possible synergistic interaction of two networks. Next, TOPS was used to print a 3D Mayan pyramid and tested under compression (**Figure 5B, Video V3**). Even after 95% strain for 3 cycles, the structure recovers back to its original shape with little deformation upon unloading. A small difference is observed in the force-strain plots between cycles 1 and 2, while little-to-no differences are observed between cycles 2 and 3, demonstrating superior shape recoverability of printed structures (**Figure 5C**). Compression tests were performed on the swelled structures, which were immersed in water for 4 days. The original structure swelled almost 5 times its volume and the compression result showed that these structures can withstand the ultimate compression strain of 84% at stress was 0.1MPa, which is 150 times smaller than the ultimate stress associated with the original structure (**Figure S14**).

**Figure 5.**
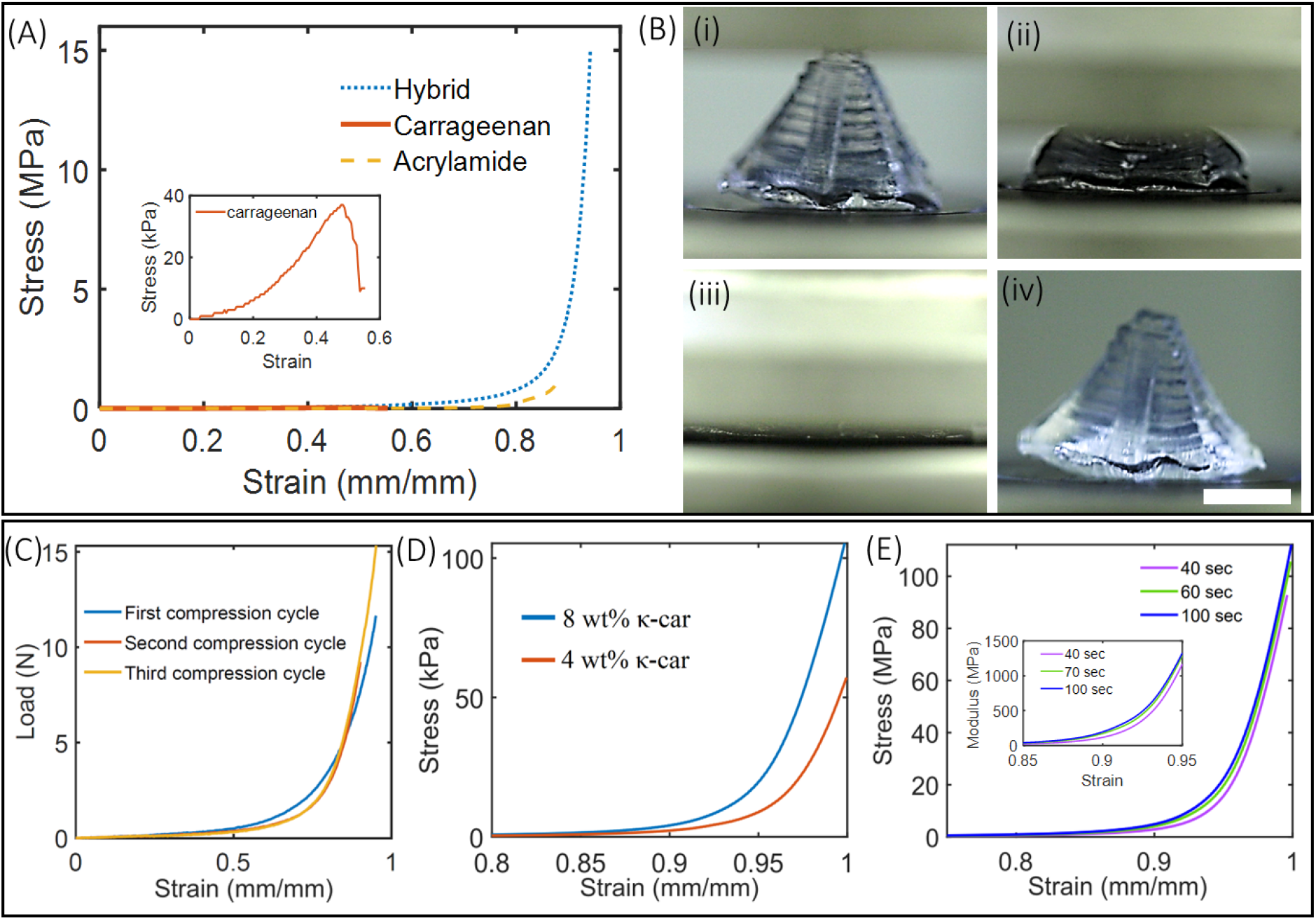
(A) Stress-strain plot obtained from cylindrical stud structure printed using acrylamide-only, κ-carrageenan only, and acrylamide/κ-carrageenan hybrid gels. Inset shows a strain-stress plot of the κ-carrageenan structure. (B) Demonstration of compression and recoverability of 3D printed Mayan pyramid structure (scale bar-5 mm). (C) Force-strain plot for 3 compression cycles of the pyramid structure. (D) Compressive stress-strain plot for structures printed using two different concentrations (4 wt% and 8 wt%) of κ-carrageenan in the hybrid gel. (E) Stress-strain plot for DN gel structures printed by varying laser exposure time at fixed laser intensity. Inset shows the change in the modulus of structures printed using different exposure times.

Specific roles of κ-carrageenan concentration and exposure times on the compressive properties of printed DN structures were also studied. Decreasing the concentration of κ-carrageenan decreases the ability of the structure to withstand loads, consistent with the trends seen in tensile testing. (**Figure 5D**). Structures printed with lower exposure times were found to softer compared to those printed using a longer exposure time (**Figure 5E, inset**). Longer exposure time strengthens the covalent bond of the acrylamide network thereby increasing the stiffness of the material.

### 2.9. Shaping DN hydrogels into a dynamically tunable soft photonic device

Here, we printed an axicon lens (a conical prism) capable of generating a dynamically reconfigurable quasi-Bessel beam and its characteristic annual ring. First, the transparency of the DN hydrogels was characterized. The transmission spectra of printed DN geometry showed a transmissivity of more than 90% over the wavelength of 400-800 nm (**Figure 6A**). A printed slab of DN gels structures clearly showed the logo of the Bioinspired Institute, depicting the high transparency of the structure (**Figure 6A, inset**). Next, TOPS was used to print an axicon lens with a laser intensity of 2.17 mW/cm^2^ and exposure time per layer of 15 seconds, and a layer thickness of 50 μm (**Figure 6B**). The diameter and thickness of the as-printed lens are 8 mm and 3.65 mm respectively and the base angle (β) is measured to be 24.5 degrees. In as-printed static conditions, a Gaussian beam passed through the axicon lens and generates an annular ring (**Figure 6C**). Then, biaxial tensile stress was applied to the DN axicon lens using a custom-built stretching device (**Figure 6D-E**). Dynamic stretching of the DN lens results in a corresponding increase in the cone apex angle and a decrease in the diameter of the annular ring as visualized using a digital SLR camera (**Figure 6F, Video V4**). Details of the optical setup and stretching devices are provided in the Methods section. Mechanically reconfigurable DN axicon lenses represent a new class of soft multifunctional photonic devices.

**Figure 6.**
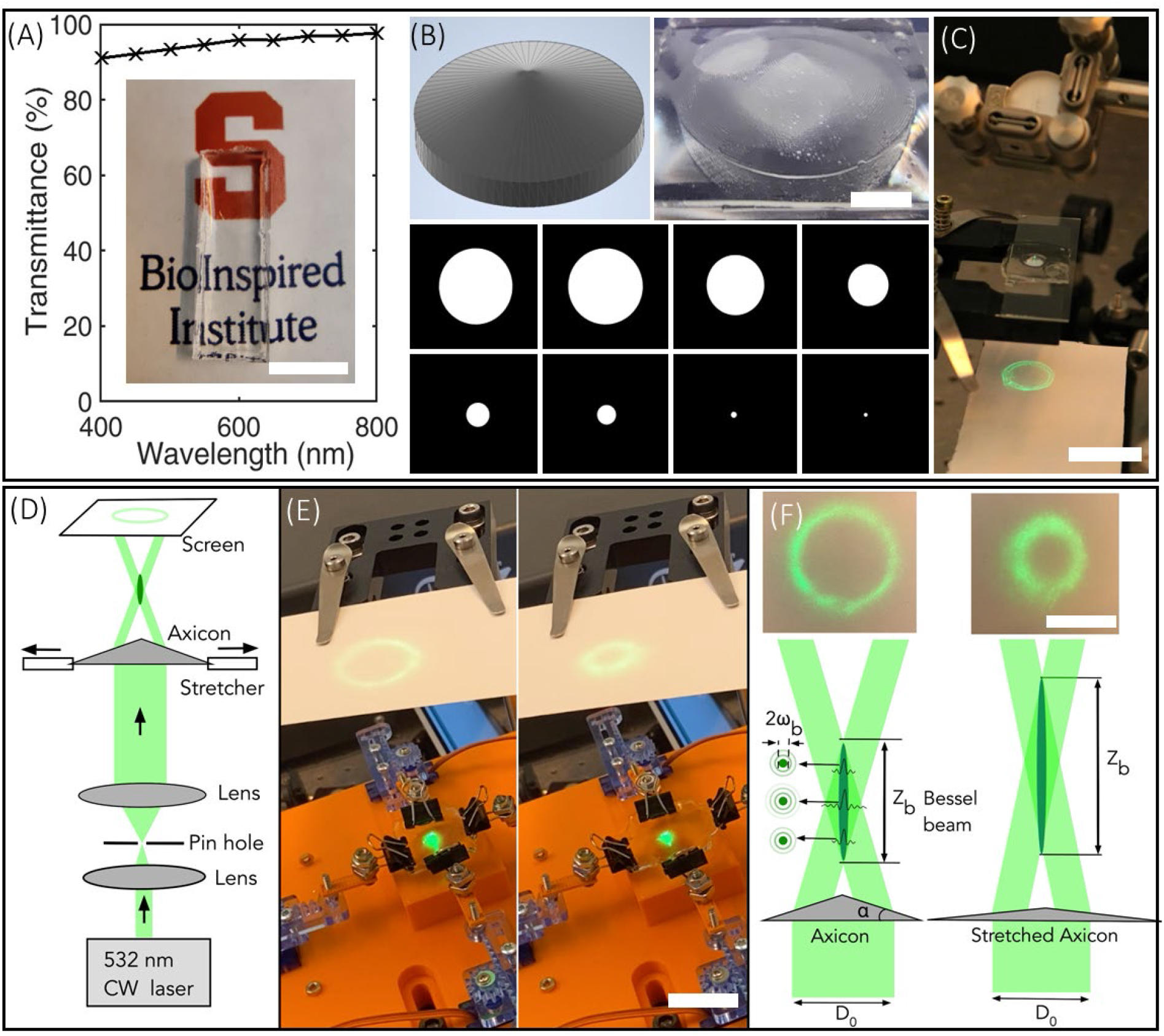
(A)Transmission spectra of structure printed using DN gel structure. The printed structure is placed on the top of the Bioinspired Institute logo to demonstrate the high transmissivity of the DN gels structure (scale bar-1 cm). (B) CAD design, computer-generated digital masks, and 3D printed axicon lens using TOPS (scale bar-2 mm). (C) Characterization of the annular ring while the axicon lens is static (scale bar-2 cm). (D) Optical setup to characterize the annular ring of the axicon lens during dynamic stretching. (E) A screenshot was obtained from Video V4 showing the tunability of the axicon lens (scale bar-3 cm). (F) Schematic showing the zero-order Bessel beam generation by the axicon and experimentally obtained annular rings before and after stretching the axicon lens (scale bar-2 cm). ω_g_=radius of the central lobe, Z_b_=length of Bessel region, D_0_=diameter of incident beam.

## 3. Discussion

This work reports TOPS-enabled DN hydrogel structures that exhibit superior mechanical properties in both tensile and compression regimes using an optimized formulation composed of acrylamide (16 wt%), κ-carrageenan (2 wt%), MBAA (0.03 wt%), and photoinitiator LAP (0.12 wt%). Photocrosslinking of the primary acrylamide network ensures the structural integrity during printing while cooling below the sol-gel transitions temperature of 80°C results in physical crosslinking of the secondary κ-carrageenan network. Since the fracture energy of the DN structures is greater than the sum of fracture energies of individual network structures, it points to the synergistic effect of crosslinking and chain entanglements between the two networks. Tensile tests show that acrylamide and κ-carrageenan networks contribute to stretchability and stiffness respectively. Results show that many processing variables modulate the mechanical properties of printed structures. For instance, the development of printed structures above 80°C and a duration of more than 3 minutes can lead to heat-induced distortions of the DN structures. An increase in crosslinker (MBAA) concentration shows low fracture energy, while an increase in exposure times increases the stiffness of the material. Necking behavior, seen in as-printed samples, can be resolved by simply immersing the samples in DI water for a few minutes which removes local defects induced by nonuniform hydration. Although possible, self-healed DN structures exhibit inferior ultimate stress and strain as compared to the as-printed structures. The compression properties of these structures are comparable to the mechanical properties of bovine cartilage and surpass previously reported 3D-printed DN hydrogel structures, as explained below.

### 3.1. Comparison of mechanical properties and resolution: State-of-art Vs TOPS-enabled DN gels structures

Among various fabrication methods at our disposal, extrusion-based direct ink writing (DIW) is the most widely used, however, achieving a resolution of 200 μm or less remains challenging.^[49]^ Multiphoton polymerization-based 3D printing techniques can print high-resolution structures at a micrometer scale, however, they are extremely slow owing to their serial point-by-point scanning.^[50,51]^ The resolution of the TOPS lithography system is comparable to different kinds of optical projection lithography such as DLP, or CLIP, which support the rapid fabrication of 3D hydrogel structures with a resolution of approximately 10-200μm. ^[34,36,39]^ (**Figure 7A**). In contrast to other work, which focuses on one regime (tensile or compression), TOPS-printed 3D structures simultaneously exhibit superior mechanical properties in both tensile regimes (strain of 2400%, stress of 130 kPa) and compression regimes (strain of 95%, and stress of 15MPa) with a high degree of recoverability. Moreover, fracture energy (1238.1 J/m^2^) and modulus of elasticity (98 kPa) are comparable to most tough hydrogels, double network hydrogels, and tough soft bio tissues. (**Figure 7B**)

**Figure 7.**
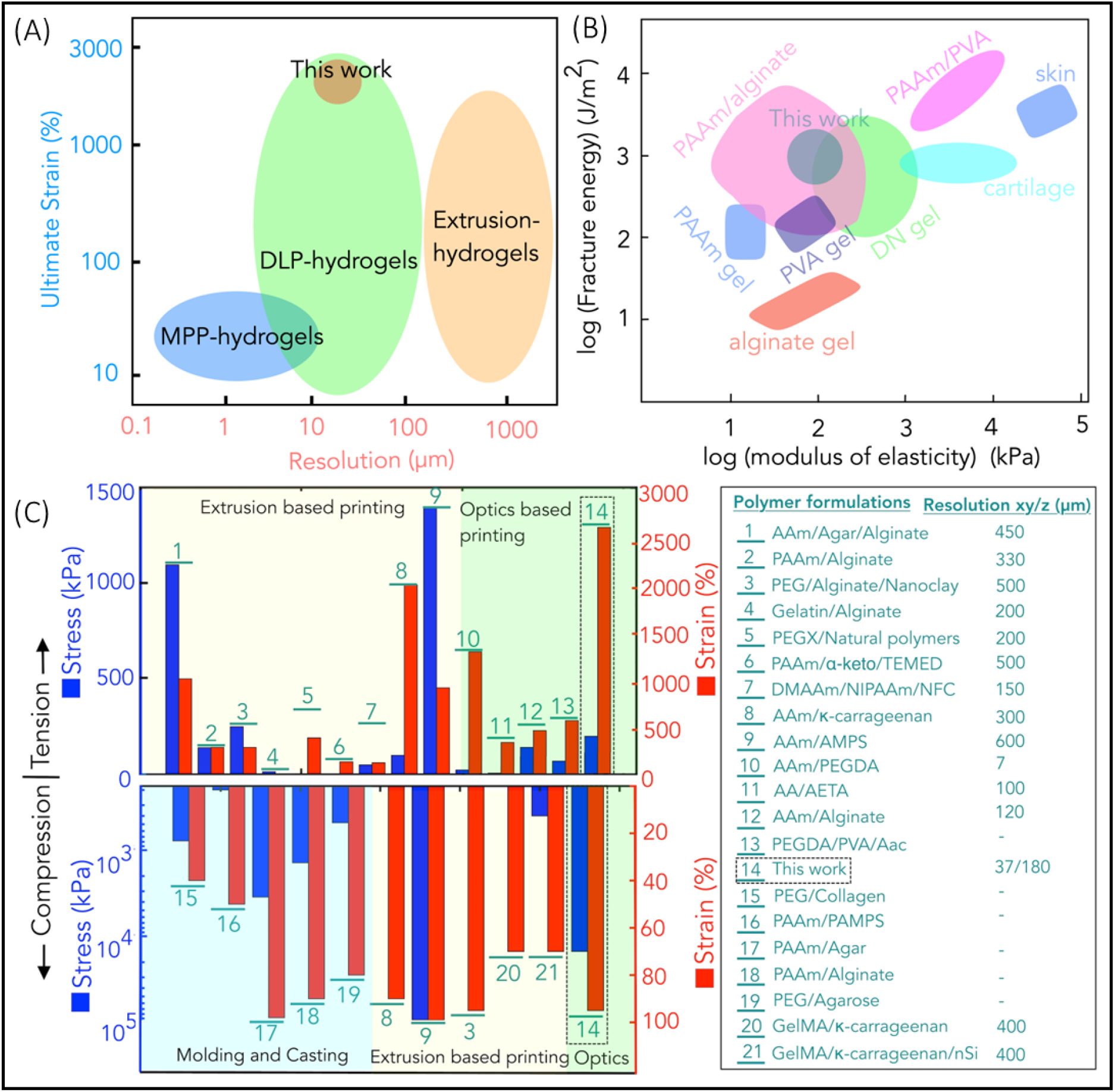
(A) Comparison of performance of TOPS printed acrylamide/κ-carrageenan structures with existing technology of hydrogel fabrication in terms of lateral resolution and ultimate strain. (B) Comparison of performance of TOPS printed acrylamide/κ-carrageenan structures with hydrogels and double network hydrogels in terms of fracture energy and modulus of elasticity. (C) Plot depicting the mechanical performance (in terms of strain and strain) and comparison of printing performance (in terms of resolution of TOPS printed acrylamide/κ-carrageenan structures with other 3D printed DN hydrogels. ***Note:** All other studies only report lateral (XY) resolution. Since we printed 3D hollow structures with overhangs and undercuts, we have also report the Z (depth) resolution* ^[49,51–70]^

As far as we know, this unique combination of high-resolution, 3D design flexibility, stretchability, compressibility, and recoverability is better than current state-of-art, which includes DN structures printed using light and extrusion-based printing methods as well as conventional casting/molding strategies. Since the literature does not report all aspects, we tried our best to compare our work with existing literature using a chart showing ultimate stress and strain in both tensile (upper part) and compression (lower part) regimes (**Figure 7C**). The best resolution was obtained for the optics-based printing of PAAm/PEGDA which showed a resolution of ~7 μm. ^[52]^ The smallest feature size obtained for our work after the development stage was 37 μm. Structure as small as 12 μm was printed, however, they did not survive the development stage. The ultimate tensile stress of TOPS DN structures is comparable with other 3D printed DN structures, whereas the ultimate strain of our samples is better than most other DN structures. Closest to our strain response in tension (2400%), extrusion-printed PAAM/ κ-carrageenan showed a strain of ~20,^[53]^ whereas the optics-based dual photocrosslinking of AAm/PEGDA showed the ultimate strain of ~12.^[52]^ Our compression strain (95%) is similar to the reported work, for instance, extrusion-based printing of PEG/Alginate/nano clay,^[54]^ molding/casting using Agar/PAAm.^[21]^ The ultimate compression stress was highest for the PAMP/AAMS, which is 93.5 MPa, however, it is not clear if the structure can recover after the compressive stress is removed. ^[55]^ In our case, a printed structure with ultimate stress of 15 MPa can recover. The only study with comparable mechanical properties was Aam/AMPS, however, the resolution of printing of this material system is limited to 600μm, while the resolution of this work is 37μm.^[55]^ Based on this, TOPS-based DN structures exhibit better mechanical properties as compared to other DN structures.

## 4. Conclusion

TOPS allowed the printing of 3D structures while maintaining the desired temperature of the prepolymer solution. This optical technique of fabrication was demonstrated to print DN hydrogel of acrylamide and κ-carrageenan at the resolution of 37 μm. The as printed 2D/3D structures were complex, mechanically strong, highly stretchable, and transparent and exhibited tunable mechanical properties by varying the printing conditions, material formulations, and post-processing steps. The printed structures performed equally well under compression and tensile force, unlike most other 3D printed DN gels structures, which only performed well under either tensile or compression force. As a proof-of-concept, a mechanically reconfigurable axicon lens was printed using TOPS, paving the way to print on-demand 3D elastomeric transparent structures can be utilized in a range of applications such as soft robotics, soft wearable electronics, adaptive optics, augmented reality, tissue engineering, and regenerative medicine.

## Supporting Information

Specifics related to materials and methods, and 15 supplementary figures, and captions for video files can be found in the supporting information document.

## Acknowledgments

The authors gratefully acknowledge Libin Yang and Prof. Zhao Qin for helping us with compression measurements and personnel in the machine shop of Syracuse University. Financial support for this project was provided by the National Institutes of Health (R21 GM141573-01) and the Syracuse University Collaboration of Unprecedented Success and Excellence (CUSE) and BioInspired Seed grant program.

