## Supplementary data for "Printing double network tough hydrogels using Temperature-Controlled Projection Stereolithography (TOPS)"

.

### 5. Experimental Section/Methods

**5.1. Chemicals.** All chemicals were used as received and were of analytical grade. Acrylamide (AAm), κ-carrageenan, N,N'-methylenebisacrylamide (MBAA), and tartrazine were purchased from Sigma-Aldrich. LAP was synthesized in our laboratory. Lithium phenyl-2,4,6-trimethyl-benzoyl phosphinate (LAP) was synthesized using a recognized method.[1] Briefly, 2.85mL dimethyl phenylphosphonite was allowed to react with 3.00mL 2,4,6-trimethylbenzoyl chloride in a covered flask for 18 hours under a nitrogen blanket with stirring. After 18 hours, 100mL of 6.25%(m/v) lithium bromide in 2-butanone solution was added to the flask. The mixture was heated to 50˚C and then allowed to react for 10 minutes. After cooling to room temperature, the white precipitate was collected via vacuum filtration and washed with four 100mL aliquots of 2-butanone. Isolated LAP was dried in a vacuum for one week to remove residual solvent.

**5.2. TOPS-based 3D printing of DN hydrogels.** TOPS-lithography optical setup used to fabricate the acrylamide/κ-carrageenan structures is shown (**Figure 1 B**). This setup consisted of a 405 nm CW laser (Toptica) which was expanded and spatially cleaned using a lens telescope and pinhole. The beam was directed toward the rotating diffuser, which changed the Gaussian intensity distribution of the laser beam to uniform intensity distribution. Rotation of the diffuser was used to average out the laser speckle. The diffuser diverged the laser beam which was collimated using a lens and then the beam was projected into the digital micromirror (DMD) device. DMD is a two-dimensional array of few micron sized mirrors that spatially pattern the laser beam, which was directed towards the projection optics by a dichroic mirror. The projection lens assembly consisted of two lenses (f=200mm), which projected the beam with unity magnification into the prepolymer solution. An imaging arm was incorporated into the setup to monitor the fabrication process. The sample holder was fabricated with copper plate and PDMS and is shown in SI (**Figure S1**). A circular disk-shaped PDMS thin film was molded and a copper plate with a circular hole (diameter 16 mm) was thermally crosslinked on top of the PDMS disk. The rim of the sample holder was prepared by molding and casting and was thermally crosslinked on the top of the copper plate. The sample holder was heated using two heating rods. A linear actuator (PI) and a controller (G910, PI) were used to control the L-shaped z-stage. A LabVIEW program was written to coordinate the switching of the DMD masks, turning the laser ON and OFF, and the z-direction movement of the stage.

**5.3. Fabrication and characterization of DN gel structure.** Fabrication was performed at an elevated temperature of 80º C. First, a custom-written MATLAB code was used to slice a 3D model of interest and was used to create a stack of the digital masks, which are binary portable network graphics (png) image files. The digital masks were uploaded to DMD, which selectively turn on/off the mirror to pattern the laser beam. This crosslinked a single layer inside the prepolymer solution. The change of the mask in the DMD was coordinated with the z-direction movement of the stage with a custom-written LabVIEW program. The laser beam photopolymerized the acrylamide to fabricate three-dimension, transparent structures. Next, the structure was placed into an ice bath or cooled at room temperature to complete physical crosslinking.

**5.4. Digital optical microscope imaging.** A Digital optical microscope (HIROX, KH-8700) was used to image and characterize the printed structure with a resolution of 1.16 µm using an MX(G)-2016(z) objective lens.

**5.5. Tensile testing.** The tensile testing was performed using a tensile tester (250lbs Actuator, Test Resources) at room temperature. Samples were printed into dog-bone shapes with a length of 6.25 mm and a gauge width of 1.50 mm. These structures were pulled at a rate of 150% strain (9.375 mm/min) using a load cell of 25 N. The modulus of elasticity was calculated as the maximum slope at the elastic region of the stress-strain plot. The fracture energy was estimated as the area under the stress-strain curve.

**5.6. Compression testing.** The compression testing was performed using a Universal testing System (Model-5966, Instron) at room temperature. Cylindrical stud structures (radius=7 mm, height=5 mm) were printed and compressed at a rate of 0.5 mm/min.

**5.7. FTIR studies.** Infrared analyses were performed in attenuated total reflection mode (ATR) using a Bruker Tensor 27 FTIR spectrometer equipped with a MIRacle ZnSe single reflection ATR block and KBr beam splitter. Spectra of the printed dual network (DN) gel samples were recorded in the range 600–4000 $cm^{-1}$ with a resolution of 4 $cm^{-1}$ and a sampling frequency of 32 scans. Samples were analyzed at three levels of hydration, both before and after tensile loading. Interfering peaks from water were removed from hydrated DN gel samples via subtraction of a spectrum averaged from replicate samples (n=9) of Nanopure deionized water (18MΩ-cm).

**5.8. Thermogravimetric Analysis.** Thermogravimetric analysis (TGA) was performed on fully dehydrated DN gel samples using a TA Instruments Hi-Res TGA 2950 (TA Instruments, Waters Corporation, Milford, MA, USA) in a nitrogen atmosphere between 25˚C and 600˚C with a heating rate of 10˚C/min. The thermal decomposition temperature was calculated at the onset of mass loss. The peak rate of mass loss was calculated using the relative maxima on a first derivative plot of weight percent vs. temperature.

**5.9. Lens stretcher.** A custom 'lens stretcher' was designed to stretch the printed optical constructs by desired increments. The stretcher was designed in Autodesk Inventor and assembled thereafter (Figure S15). The base plate was 3D printed and designed to affix to a Thorlabs optical fixture. This allowed the entire stretcher to be mounted in line with the light path. The 3D-printed axicon lens is placed on the raised center platform and the central hole in the base plate allows light to pass through both the plate and the lens. Four servo motors with linear actuation gearing were used to achieve uniform stretching of the constructs from all four sides. The servo motors were programmed via Arduino to move uniformly to any desired position in their range of motion. Small binder clips were fastened to the linear slides so the constructs could be attached firmly without slipping or tearing.

### Dimension of the sample holder

The dimensions associated with the geometry are provided in Table 1.

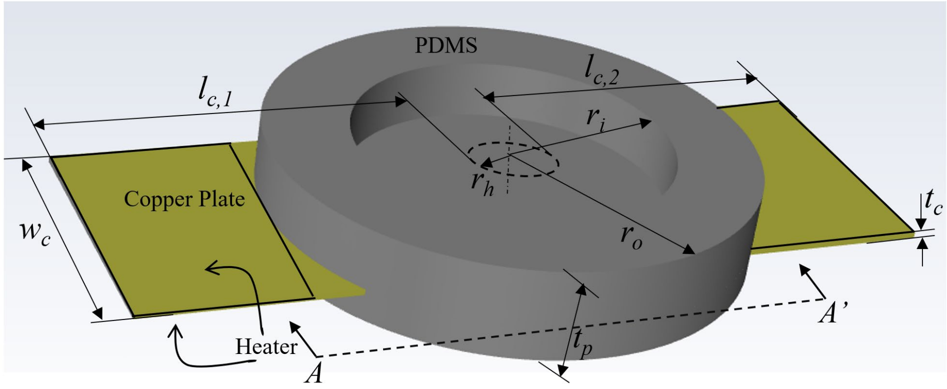


**Figure S1**. Computational Domain consisting of PDMS dish connected with copper plate.

*Table S1. Dimensions of geometry associated with the computational domain are shown in Fig.1.*

| Symbol | $w_c$ | $l_{c,1}$ | $l_{c,2}$ | $r_i$ | $r_o$ | $r_h$ | $t_p$ | $t_c$ |
|---|---|---|---|---|---|---|---|---|
| Dimension (mm) | 53 | 65 | 70 | 30 | 38 | 8 | 17.5 | 1.1 |

### Heat Distribution Simulation

It was important to maintain a critical temperature of the solution throughout the fabrication process, hence we designed a CAD model of the sample holder and studied the temperature distribution using simulations. The design of the sample holder consisted of a copper plate with a center hole embedded inside the PDMS bath (**Figure S1**). Two heated rods on either side of the dish were designed to heat the copper plate and the 16mm diameter hole in the Cu plate acted as the fabrication window. To gain a better insight into the temperature distribution over the PDMS layer of the sample holder design, we performed computational fluid dynamics (CFD) simulation with conjugate heat transfer. The computational domain consisted of the designed PDMS dish and a copper plate extended on either side of the dish (**Figure S1**). At the top and bottom surface of the copper plate, a constant temperature boundary condition with T = 416 K (142.85°C) was applied to mimic the heater (used in an experimental study). All other surfaces of the geometry were provided with convection heat transfer to ambient temperature ($T_{amb}$ = 300 K (26.85°C).

The simulation was performed by discretizing the computational domain into a finite number of control volumes (or grid points) and by simultaneously solving the physical equations of continuity, momentum, and energy, in each point to obtain the spatial and temporal distribution of temperature. For the computational domain, a mesh with 160,000 grid points was utilized to obtain the temperature distribution (**Figure S2**). The distribution was obtained for various time points until the steady state is attained. After the simulation study, the mesh was refined, and the simulation was repeated for meshes with 200,000, 240,000, and 280,000 grid points to investigate the grid sensitivity of the result. By comparing temperature distribution over the PDMS layer for different meshes, one with 240,000 grid points was found to be an optimum mesh which is utilized for further study here.

Finite volume methods-based commercial solver ANSYS Fluent was utilized to solve the equations. In the simulation, the entire domain was first initialized with $T_{init}$ = 353 K (79.85 °C), and the ambient was set at 300 K (26.85°C). The operating pressure was 1 atm. **Figure S3** shows the temperature distribution over the PDMS layer at different time instants obtained using the optimum mesh. At approximately t = 90 s, the spatial distribution of temperature attains a steady state and further continuing the simulation shows no changes in the state.

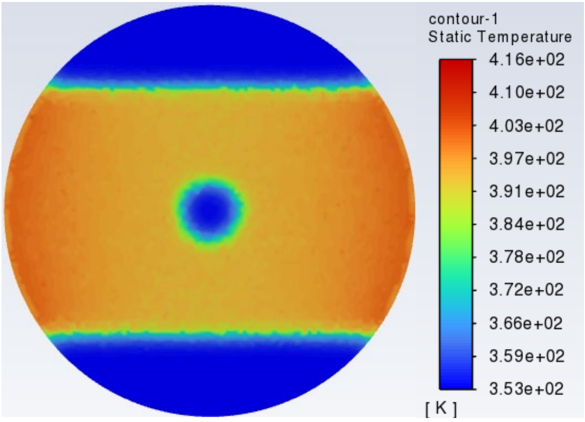


**Figure S2.** Temperature distribution over the PDMS layer at the plane corresponding to sections A-A' shown in Figure S1.

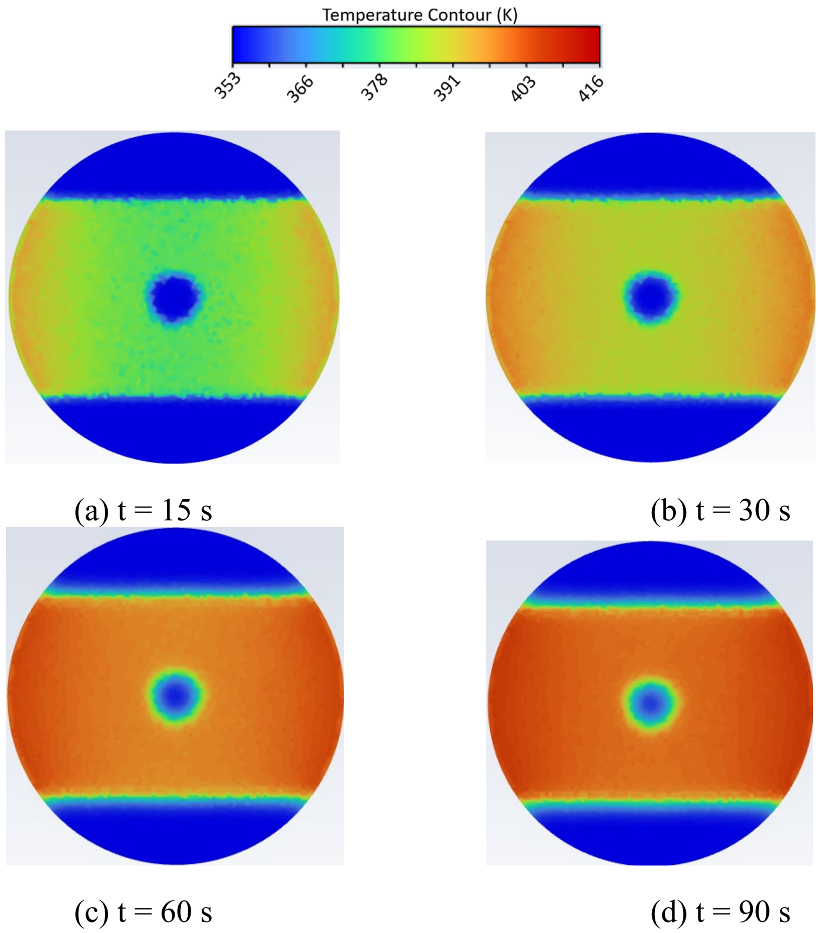


**Figure S3.** Temporal evolution of temperature distribution over the PDMS layer heated using the copper heater. The plots are obtained from the simulation using an optimum mesh of 240,000 grid points.

**Effect of exposure time on the tensile properties of TOPS printed DN gel dog-bone structures**

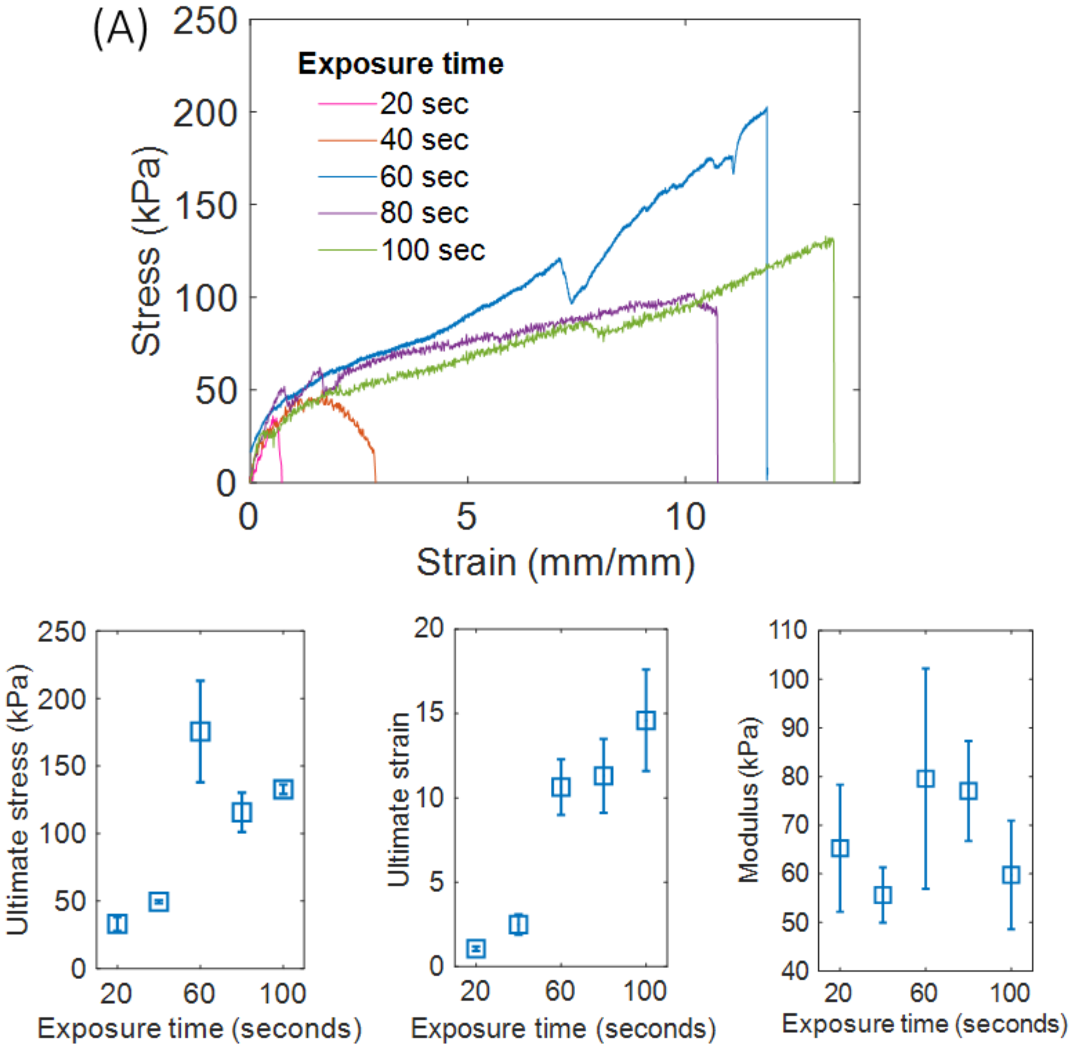


**Figure S4.** (A) Stress−strain plots obtained from TOPS printed DN gel structures exposed to different exposure times. Ultimate stress, ultimate strain, and tensile modulus recorded for the TOPS printed structures with varying exposure times. (Error bars: Mean±SD)

**Effect of composition of photoinitiator on the tensile properties of TOPS printed DN gel dog-bone structures**

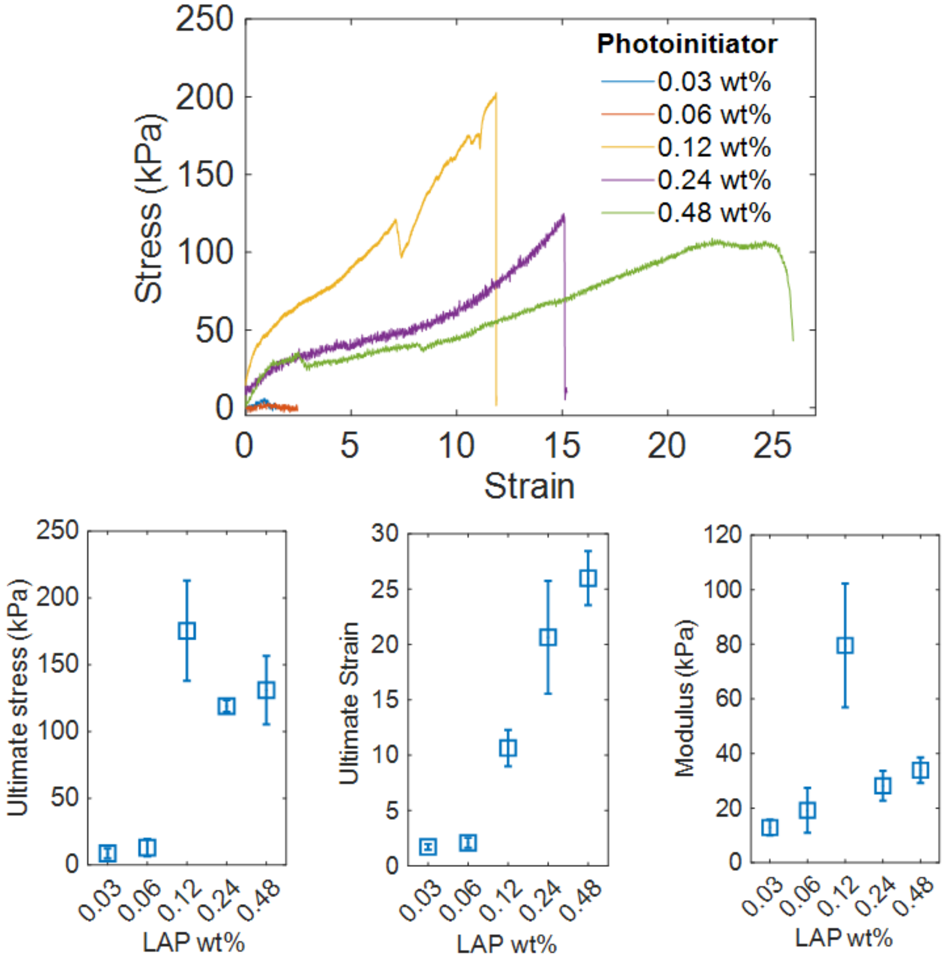


**Figure S5.** Stress-strain plot obtained from the TOPS printed hybrid gel structure with varying proportions of photoinitiator. Ultimate stress, ultimate strain, and elastic moduli obtained for the DN gel structures with varying concentrations of photoinitiator. (Error bars: Mean±SD)

**Effect of composition of crosslinker on the tensile properties of TOPS printed DN gel dog-bone structures**

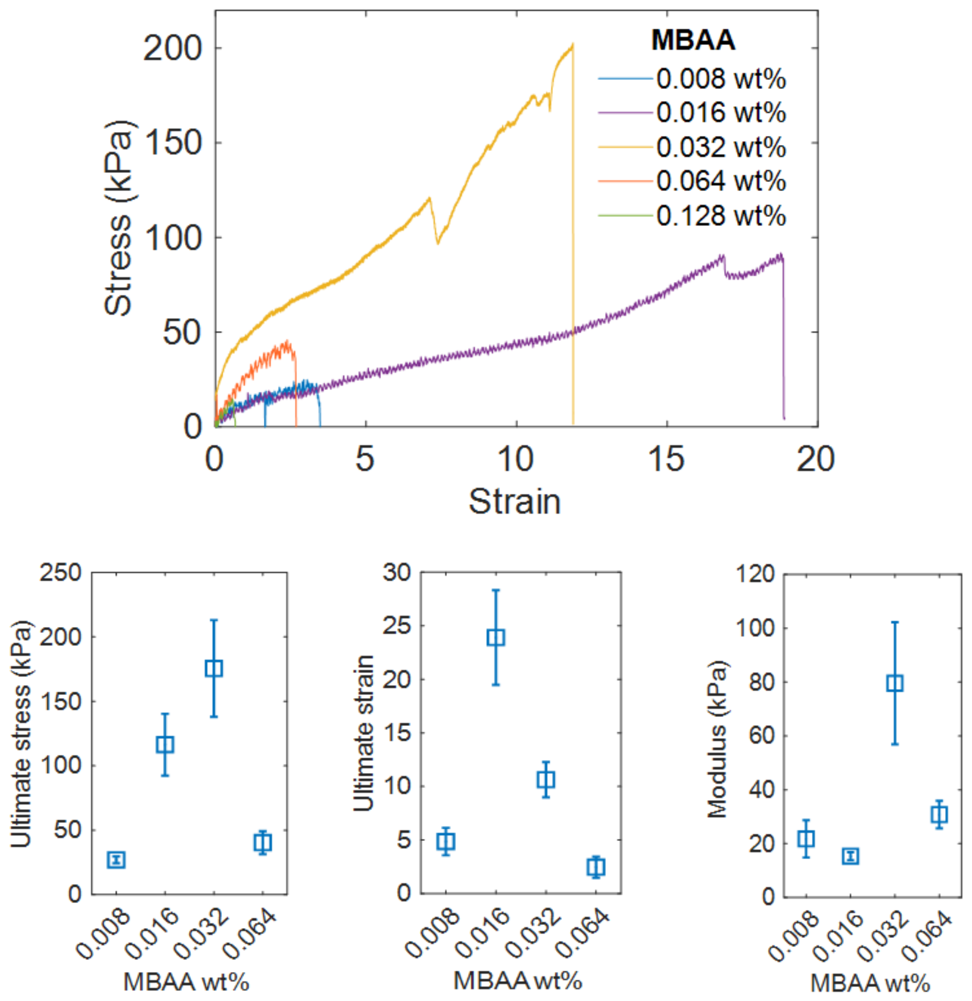


**Figure S6.** Stress-strain plots obtained from the TOPS printed hybrid gel structures printed with varying proportions of MBAA crosslinker (0.008 wt%, 0.016 wt%, 0.032 wt%, 0.064 wt%, and 0.128 wt%,). Ultimate stress, ultimate strain, and elastic moduli for the structures printed with varying amounts of MBAA crosslinker. (Error bars: Mean±SD)

**Effect of composition of κ-carrageenan on the tensile properties of TOPS printed DN gel dog-bone structures**

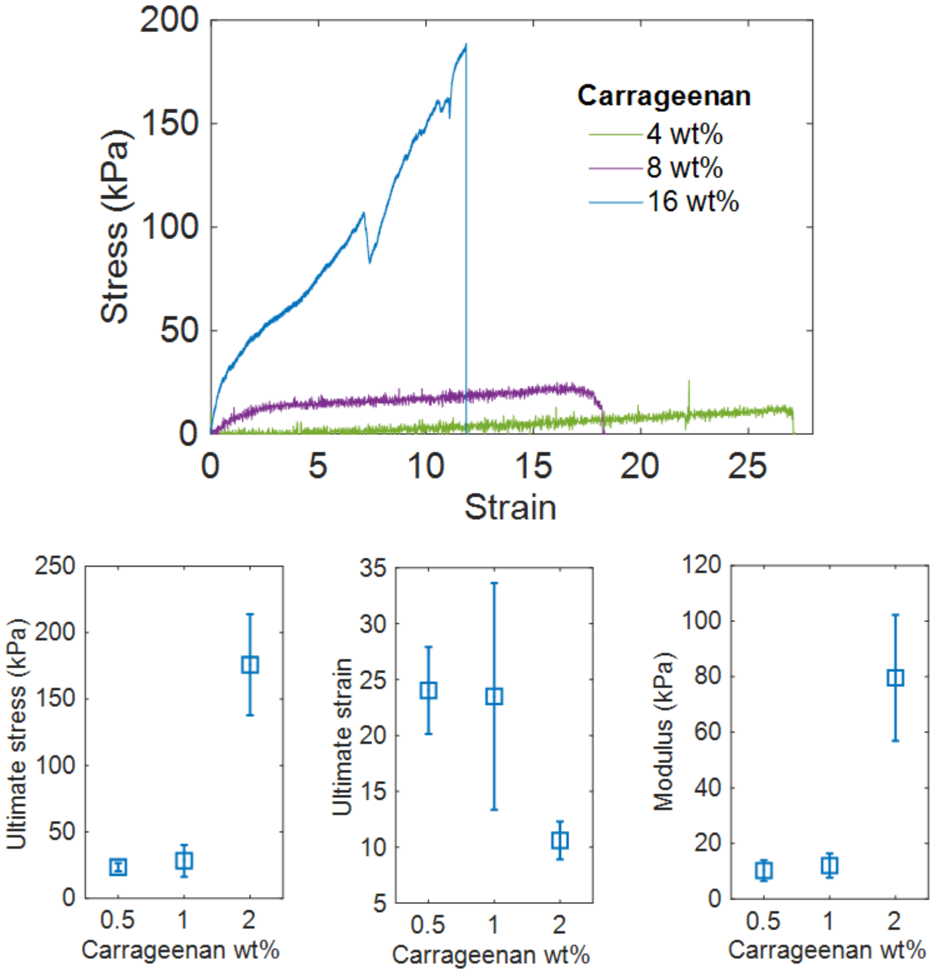


**Figure S7.** Stress-strain plots obtained from the hybrid structures printed with varying proportions of κ-carrageenan (0.5 wt%, 1 wt%, and 2 wt%). (Error bars: Mean±SD)

### Swelling of DN gel structures

DN cylindrical stubs printed via TOPS were immersed in DI water. Results show that stub diameters and heights increased by 21% during the first hour, reached 45% in one day, and saturated after 78 hrs (**Figure S8**). In terms of mass, the printed structure (right after the printing) weighed 0.5 gm and the structure absorbed 7.9 gm of water, which is 17 times the original mass. Total water content before swelling was 81% and this increased to 98.8% after immersing the structure in water for 78 hours.

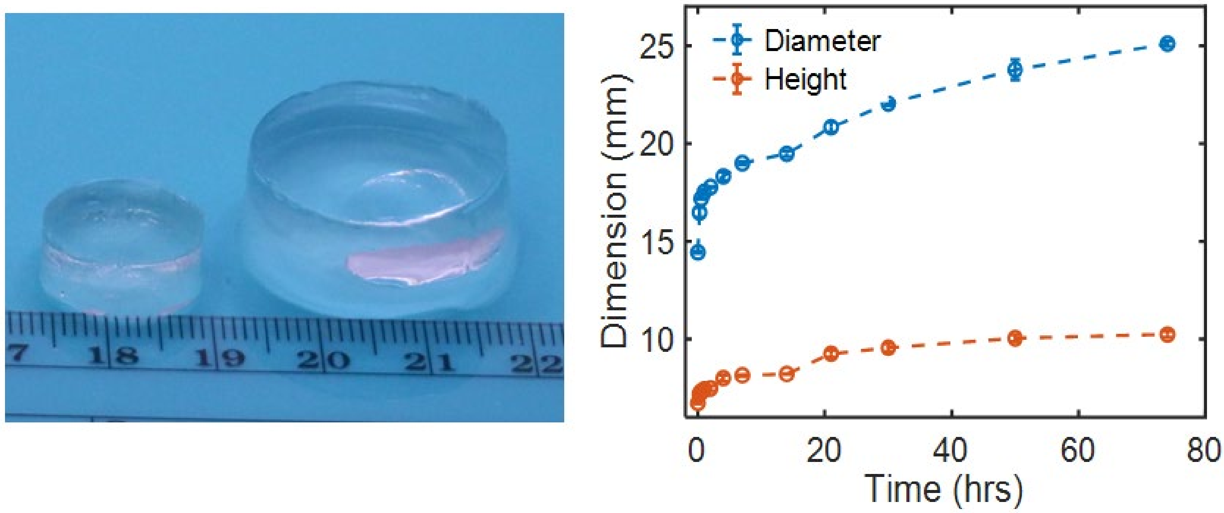


**Figure S8.** Image and plot of swelling of TOPS printed DN gels cylindrical stud and associated dimension (Diameter and Height) recorded for 78 hours.

### Influence of swelling on tensile and compression properties

Tensile properties of DN dogbone samples swollen for 5 min, 10 min, 4hrs, and 4 days were studied. The 4-day swollen sample was too soft to reliably handle, and hence it was omitted from this study. Results show that the ultimate stress, the ultimate strain, and the modulus were highest for samples swelled for 5 minutes as compared to samples swelled for 10 minutes and 4 hours.

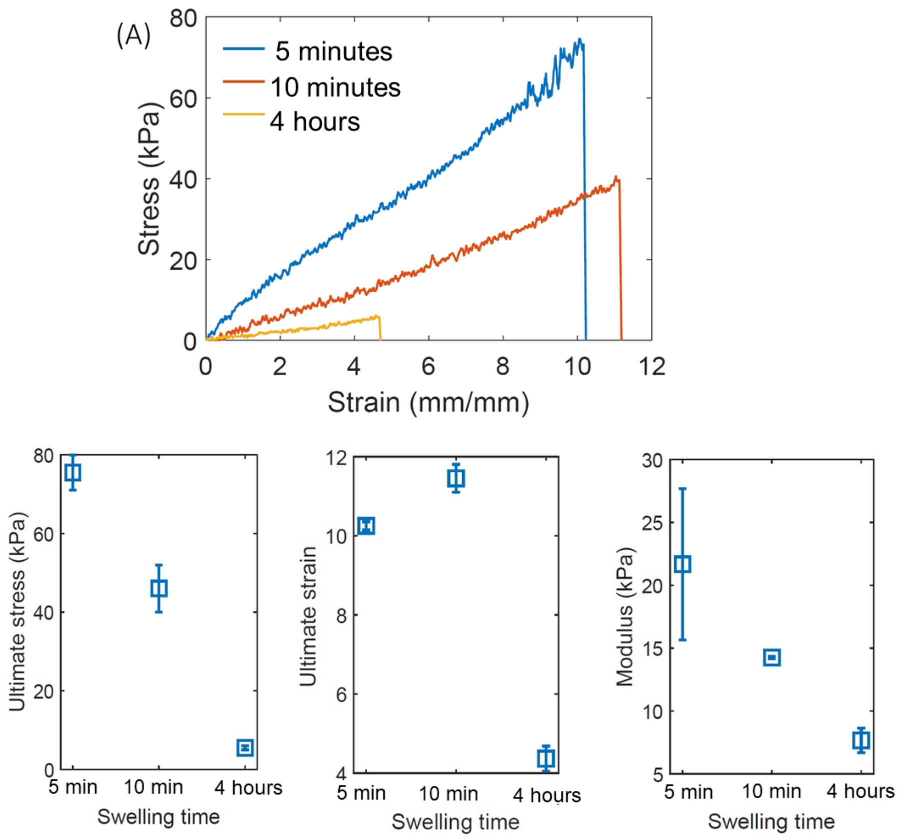


**Figure S9.** (A) Stress-strain curves (B) Stress-position curves recorded from the hybrid gel structures swelled for 5 minutes, 10 minutes, and 4 hours. Ultimate stress, ultimate strain, position, and elastic moduli for the hybrid structures swelled for different time points. (Error bars: Mean±SD)

**Effect of exposure time on the tensile properties of swelled DN gel structure**

Longer exposure time during TOPS printing resulted in lesser swelling and therefore exhibited better mechanical properties. For instance, structures exposed for 2 minutes swelled 1.28 times of original length in 4 hours, whereas the structure exposed for 1 minute swelled 1.85 times its length at the same time. Longer light exposure during TOPS, when swelled for 4 hours withstood the larger ultimate stress of 14±2 kPa and modulus 14.95±3.7 kPa. These parameters were 5.5±0.5 kPa and 7.68±0.9 kPa for the structure exposed for 1 minute. Further, the strain was almost double (8.05±1.45) for longer exposed structures (**Figure S10**).

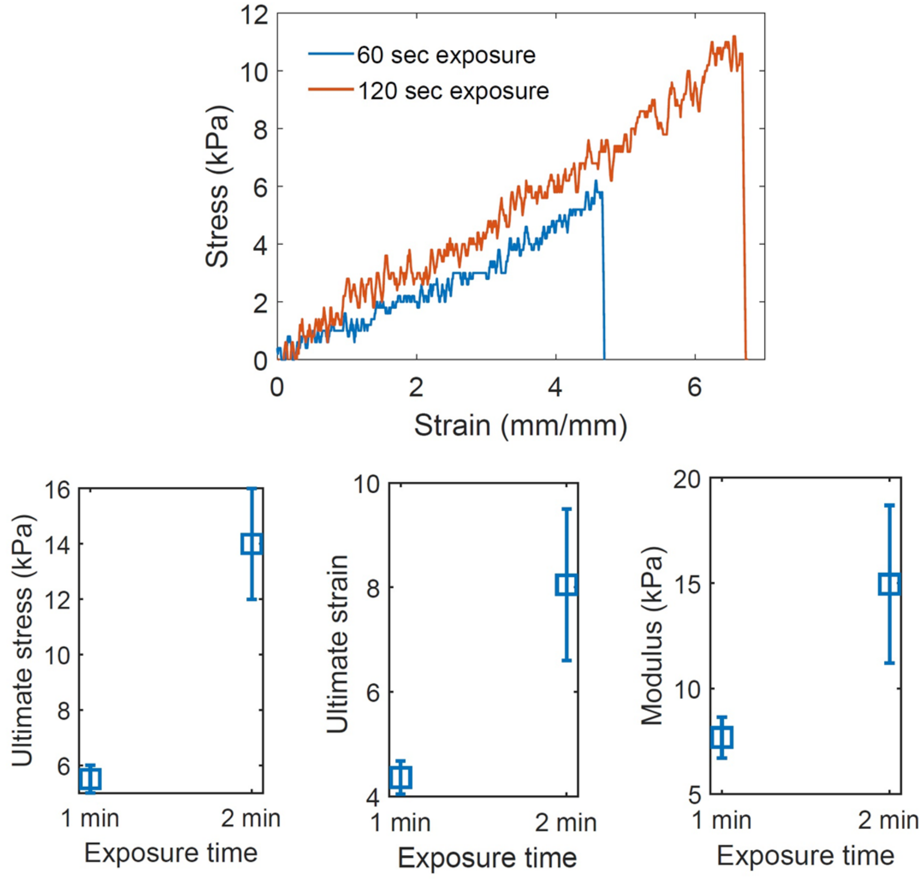


**Figure S10.** Stress-strain plots obtained from the swelled structures printed using different exposure times (60 seconds and 120 seconds). Structures were swelled for 4 hours in water. Ultimate stress, ultimate strain, position, and elastic moduli obtained from swelled structures printed using different exposure times.

**Effects of hydration, dehydration, and rehydration**

The ability of the printed structure to recover after dehydration followed by rehydration was tested. As-printed DN dogbone structure, dried for 2 days using a dehumidifier, was rehydrated in water for 30 mins until the size reaches 1.3 times the size of the as-printed structure. Tensile tests showed that these samples regained their ultimate stress (24.5±1.5 kPa), ultimate strain (7.675±0.205). Similar results were obtained when the as-printed samples were first hydrated completely, dehydrated completely, and rehydrated to 1.3 times the size of as-printed samples with ultimate stress (24±0 kPa), ultimate strain (9±0.6), and modulus (11.11±0.12 kPa) (**Figure S11**).

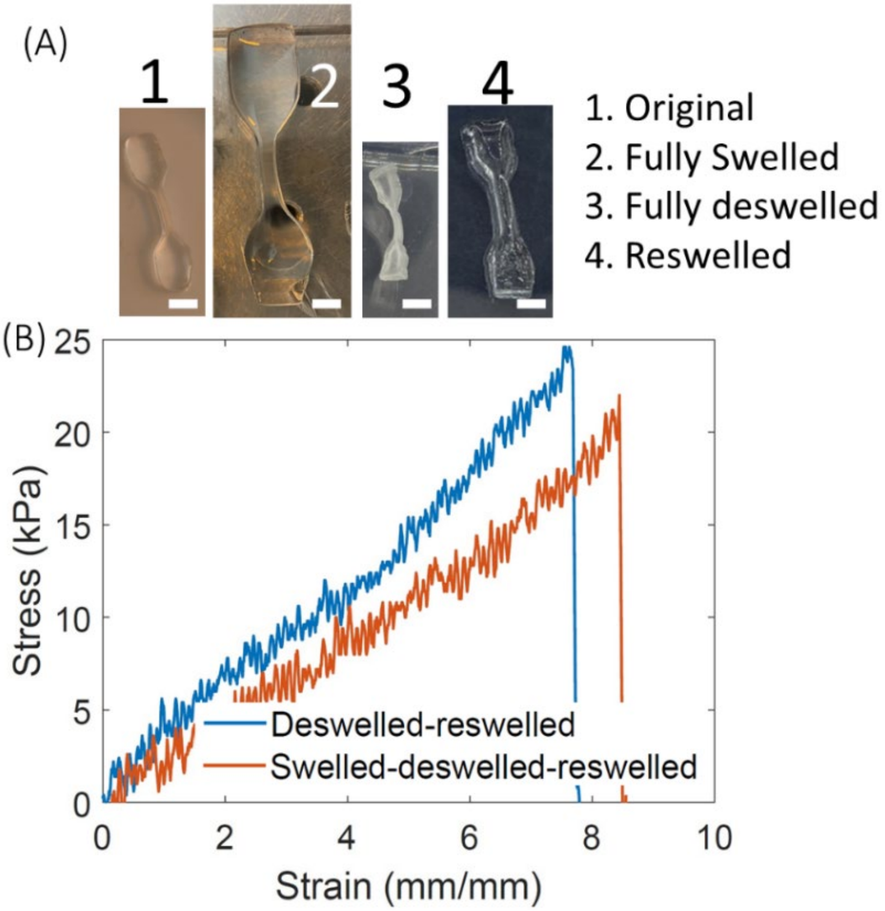


**Figure S11.** (A) Images of dog-bone structures after swelling and deswelling. (B) Stress-strain plots were obtained from the structures after different stages of swelling and deswelling. (Scale bar= 5mm)

*Table S2. Assignment of Relevant FTIR fingerprints associated with functional groups obtained from Figure 4C(i).*

| Wavenumber ($cm^{-1}$), PAAm | Wavenumber ($cm^{-1}$), DN gel (Tensile Unloaded) | Assignment |
|---|---|---|
| 1435 | 1437 | ν, C─N |
| 1454 | 1456 | δ, C─H |
| 1600 | 1598 | δ, NH2 (amide II) |
| 1668 | 1672 | ν, C═O (amide I) |

### Self-healing of the 3D-printed structures

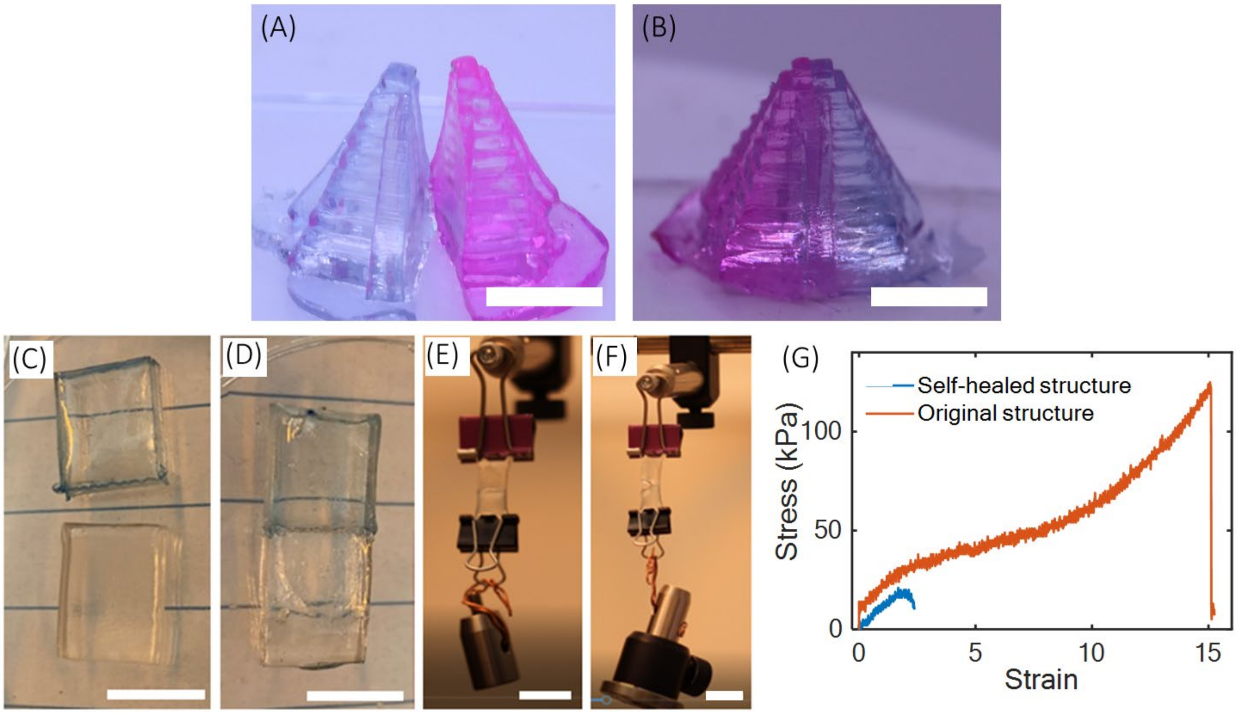


**Figure S12.** (A,B) Demonstration of self-healing properties of 3D printed Mayan pyramid structure (Scale bar- 5 mm). (C,D) Self-healing of rectangular slab printed using acrylamide/-carrageenan structure (scale bar- 10 mm). (E,F) Performance of self-healed structure subjected to a load of (70 gm) and (200 gm) (Scale bar-10mm) (G) stress-strain plot comparing the tensile performance of a self-healed structure with that of the original structure.

**Thermogravimetric analysis (TGA) studies on DN gels structures**

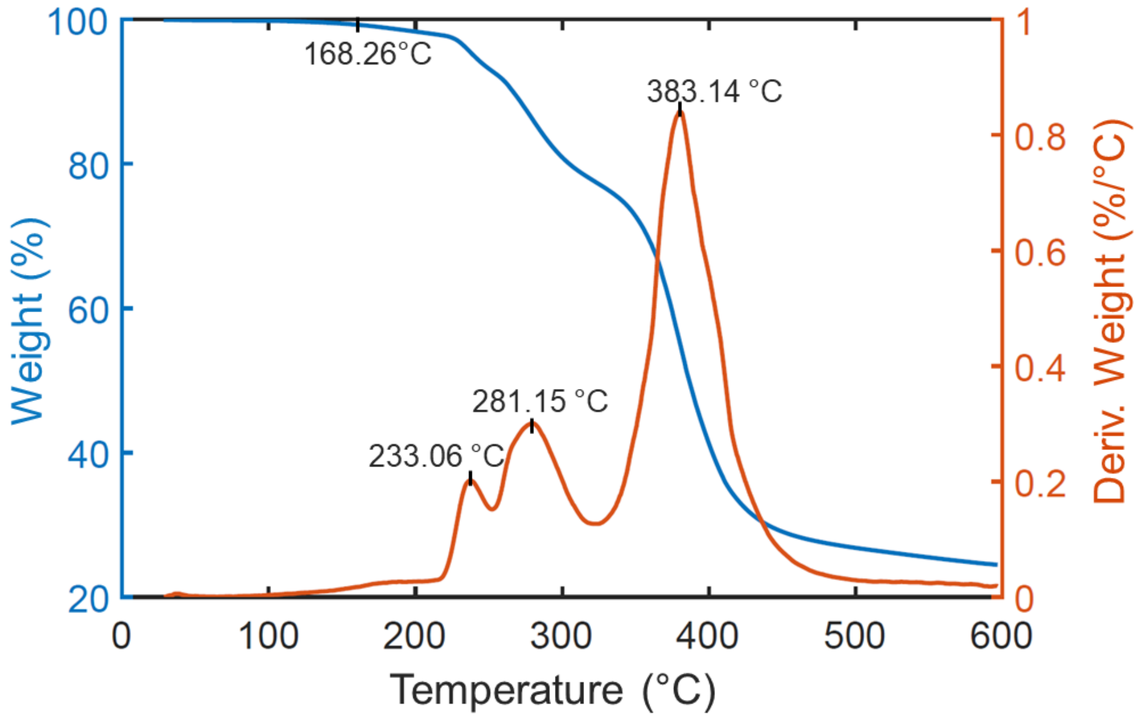


**Figure S13.** TGA plot obtained from fully dehydrated DN gel under a nitrogen atmosphere between 25˚C and 600˚C with a heating rate of 10˚C/min.

**Effect of swelling on the compressive properties of DN gel structures**

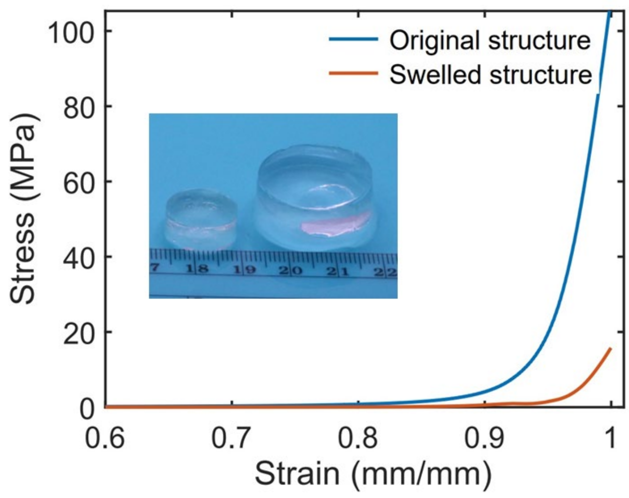


**Figure S14.** Stress-strain plot was obtained by compressing the original structure and the structure swelled for 4 days. Inset shows the original structure and swelled structure.

### Lens stretcher

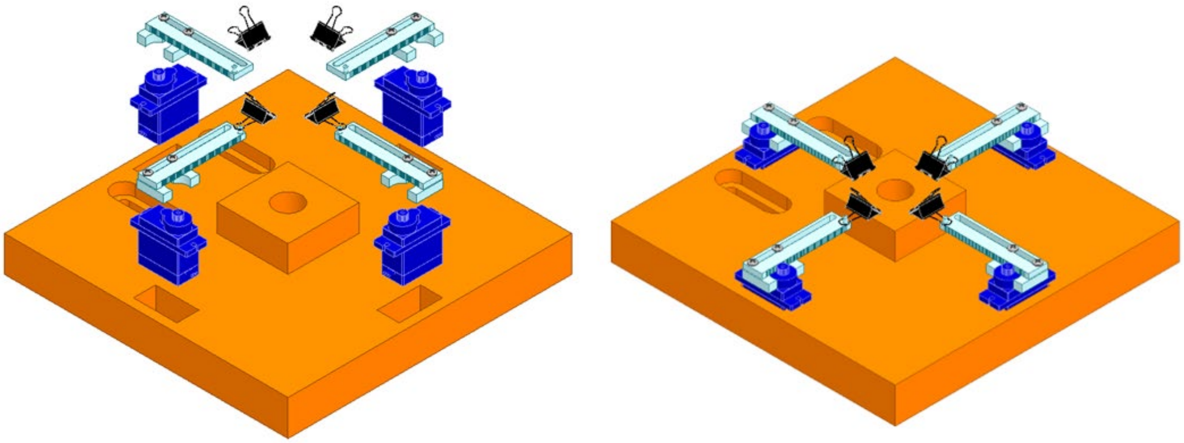

**Figure S15.** CAD design of before and after assembly of axicon lens stretching device.


### Legends of supplementary movies

***Movie V1.*** *Stretching of TOPS printed acrylamide/κ-carrageenan double network hydrogel dog-bone structure. This movie shows the stretching of structures with necking and without necking.*

***Movie V2.*** *Stretching of TOPS printed acrylamide/κ-carrageenan double network hydrogel 3D structure.*

***Movie V3.*** *Compression of TOPS printed Mayan Pyramid using acrylamide/κ-carrageenan double network hydrogel. This movie shows that the structure regains its structure after compression force is lifted.*

***Movie V4.*** *Tunability of the axicon lens using a custom-designed lens stretcher. This movie shows a change in the size annular rings of the Bessel beam when the lens is stretched.*
